# Persist or Give up? Fire ants motivated to search for a high-quality food source even if they don’t know how to find it

**DOI:** 10.64898/2025.12.23.696297

**Authors:** Chinmay Hemant Joshi, Anna Dornhaus

## Abstract

Finding resources for the colony is one of the most difficult and risky tasks for a social insect worker. A worker on a foraging trip can face a number of challenges, including interference from other individuals, her own errors, and environmental disturbances. Collectively, colonies may use a variety of strategies to minimize the impact of such perturbations on the foraging process. Here, we investigated how individual *Solenopsis xyloni* ant workers react to perturbation of an established pheromone trail. We trained foragers from colonies in the field to either a low or high concentration sucrose solution in a feeder on a T-maze setup, then replaced a section of floor covering, removing a section of the pheromone trail previously laid. We found that while ants made correct choices on the T-maze when the trail was intact, their choices did not differ from chance when the trail was absent, indicating strong reliance on a pheromone trail (and not, for example, memory) to return to the resource. Moreover, when the trail was absent, we found that a majority of ants abandoned the resource, and that even the ants that were able to reach the resource did not repair the perturbed trail. However, with a high-quality resource, more ants persisted in attempting to reach it (instead of abandoning). We interpret these responses in the framework of robustness mechanisms discussed in systems biology. Our study thus links individual and collective responses to perturbations, and provides an empirical example of how information use interacts with system robustness.

**Statements and declarations:** The authors have no competing interests to declare that are relevant to the content of this article.

## Introduction

Social insects are complex adaptive systems that perform various tasks, such as foraging cooperatively (Bonabeau 1998; Garnier et al. 2007). Worker behavior has evolved to maximize the collective foraging performance of the colony (Wilson 1971; Garnier et al. 2007; Doering et al. 2024). Individual workers in a colony can use a variety of ways to communicate information about food sources to their nestmates, often leading to recruitment towards the food source (Frisch 1967; Hölldobler and Wilson 1990; Nieh 1998; Czaczkes et al. 2015; Leonhardt et al. 2016; Ganeshaiah 2023). These communication systems differ in several aspects, including signal modality, information content, how signals are reinforced to sustain recruitment, and how they are adapted to the local environmental conditions (e.g., Lanan 2014; Leonhardt et al. 2016). However, communication systems can be affected by sudden changes (i.e., perturbations) in the environment. Perturbations such as fluctuations in air pressure (Sujimoto et al. 2020), rain (Farji-Brener et al. 2018), surface temperature (van Oudenhove et al. 2011), and wind (Alma et al. 2016) can affect communication among workers and thereby collective foraging performance. Specifically, these perturbations can severely affect workers directly (Alma et al. 2016; Farré-Armengol et al. 2016; Farji-Brener et al. 2018), affect their communication signals (van Oudenhove et al. 2011, 2012; Alma et al. 2022) or ability to amplify these signals to sustain recruitment (van Oudenhove et al. 2012; Zhang et al. 2020), or affect the food source itself (Farji-Brener et al. 2018).

Robustness is an emergent, system-level property that allows a system, such as an ant colony, to maintain its performance when subjected to different perturbations (Kitano 2004; Masel and Siegal 2009). Robustness arises from how its components interact with each other and with the environment around them (Kitano 2004, 2007; Wagner et al. 2007; Whitacre 2010; Nijhout et al. 2019). Kitano (2004), in his seminal conceptual review, proposed the following main mechanism categories, which are thought to apply across a variety of complex systems: (1) ***feedback loops***, (2) ***buffering***, (3) ***fail-safes***, and (4) ***modularity. Feedback loops*** (or ‘system control’) allow the system to return to its original state by counteracting the perturbation (negative feedback loop) or by moving the system to another, more suitable state given the new environment (positive feedback loops). ***Buffers*** (or ‘decoupling’) entail having more capacity in existing functional units than normally needed, so that the system can withstand a perturbation. ***Fail-safes*** (or ‘redundancy and diversity’) involve the existence of additional functional units that can take over for any failed components. Finally, ***modularity*** entails having separate (yet interconnected) modules that allow the system to restrict negative effects to a particular module to prevent their spread to other modules. These mechanisms have been identified and studied primarily in genetic networks and biochemical pathways across different organisms (Alon et al. 1999; Freeman 2000; Conant and Wagner 2004; Fritsche-Guenther et al. 2011; Holme 2011), within engineered systems (Tell et al. 2022; Kiakojouri et al. 2023), and in designed (human) organizational frameworks (Xie et al. 2022; Duvald et al. 2025).

Both molecular systems and social insect colonies contain individual components whose actions and behaviors have been optimized at the collective level (Seeley 2002; Kitano 2004; Garnier et al. 2007). Therefore, the systems-level thinking developed in molecular systems can potentially be applied to understand how social insects respond to different perturbations (and vice versa to study molecular systems). Social insects use various strategies to maintain foraging performance that fall under one or more of the above categories. Workers may use inhibitory signals or modulate positive communication to steer nestmates away from exhausted and/or suboptimal food sources (Evison et al. 2008; Czaczkes and Heinze 2015; Alma and Buteler 2019) - both examples of feedback loops maintaining a kind of homeostasis in foraging. Alternatively, workers can flexibly alter their behavior to continue foraging amidst perturbations (strong wind gust, for instance) (Gordon et al. 2013; Sujimoto et al. 2020; Alma et al. 2022), an example of decoupling. Sampling multiple sources of the same signal (Tanner and Visscher 2008) or combining information from multiple information sources (Czaczkes et al., 2011, 2013; Letendre & Moses, 2013) could likewise be seen as an example of buffering (more capacity available than typically needed). If insects on the other hand switch between information sources when one becomes unreliable, this could be considered a fail-safe mechanism: such as when bees switch to using their memory when social signals are unreliable (Grüter et al. 2011; Dunlap et al. 2016; Jones et al. 2019).

Our aim in this study was to identify mechanisms used by social insects to maintain their collective performance in the face of environmental perturbations. Specifically, we focus on how ants deal with the disruption of communication during foraging, and which of the categories of robustness mechanisms may be used.We used southern fire ants (*Solenopsis xyloni*) as a model system for this study, and disrupted their foraging by removing a section of their pheromone trail to a food source. We then examine the responses of individual ants. We test the following specific hypotheses, which are matched to the types of robustness mechanisms proposed by Kitano 2004: (1) Ants use a fail-safe mechanism, i.e., an alternative mechanism to locate the food source, such as memory, (2) Ants employ a buffer mechanism in the face of pheromone trail disappearance, such as continuing to walk in the same direction as the original trail to relocate the food source, (3) A disrupted trail causes negative feedback (i.e., compensatory behavior), such as increased deposition of trail pheromone on the path to food, (4) Ants abandon the perturbed trail to redirect towards another food source nearby, and (5) Finally, it is also possible that ants show little robustness to disruption of their trail communication, and that foraging performance is severely impacted. We use a classic T-maze setup to examine these hypotheses, and a food source of low or high value to test whether increased food source value results in different strategies used by ants.

## Materials and methods

### Study area and Study species

The southern fire ant (*Solenopsis xyloni*, McCook (Myrmicinae)) is one of the most common ant species across the Southwestern U.S (MacKay and Mackay 2002). *Solenopsis xyloni* nests in soil or under stones in open areas and near the edges of lawns in urban and semi-urban areas (Trager 1991; MacKay and Mackay 2002). *Solenopsis xyloni* relies upon trail pheromones to recruit nestmates to food sources (Barlin et al. 1976; Tschinkel 2013). This pheromone is produced in the Dufour’s gland and extruded via their sting, which is both connected to the Dufour’s and venom glands (Barlin et al. 1976). Individual fire ants press the stinger against the ground and drag it on their way to the nest from a food source (Hangartner, 1969, and personal observations in *S. xyloni*). We conducted this study on wild colonies at the University of Arizona campus (Tucson, Arizona, 32.23^°^N, 110.95^°^S). In total, we used 17 ant colonies across four years (July-November, 2022-2025).

### Experimental setup

We used a laser-cut plexiglass T-maze (arm length = 12.6 cm, stem length = 8.6 cm, thickness = 0.32 cm) supported by cardboard columns (diameter = 0.80 cm) attached to small Petri dishes (diameter=3.3 cm) (Fig. 1a). To ensure that ants only entered and exited through one point in the setup, we applied Fluon (Company name: byFormica) on the underside of the T-maze and the ramp, and filled the petri dishes with water. The removable ramp allowed us to restrict the entry of ants during the experimental trials (such that we could isolate a single focal ant, see below). We cut printer paper to the dimensions of the ‘T’ of the maze, and cut it into sections so it could be selectively removed (Fig. 1b, and see below for more details). We attached these paper sections as surface covering using washable glue (Elmer’s Repositionable Washable Glue Stick). Lines were lightly drawn on the paper with black pencil to mark key measurement locations (Fig. 2a). We used an Eppendorf tube (1.5 ml) as a feeder by punching small holes on the side near the cap, and attaching it upside down to the maze with a small lump of putty to stabilize the feeder (Fig. 1b). The feeder was filled with a sucrose in water solution (either 1.5 M or 3 M sucrose).

**Fig. 1.**
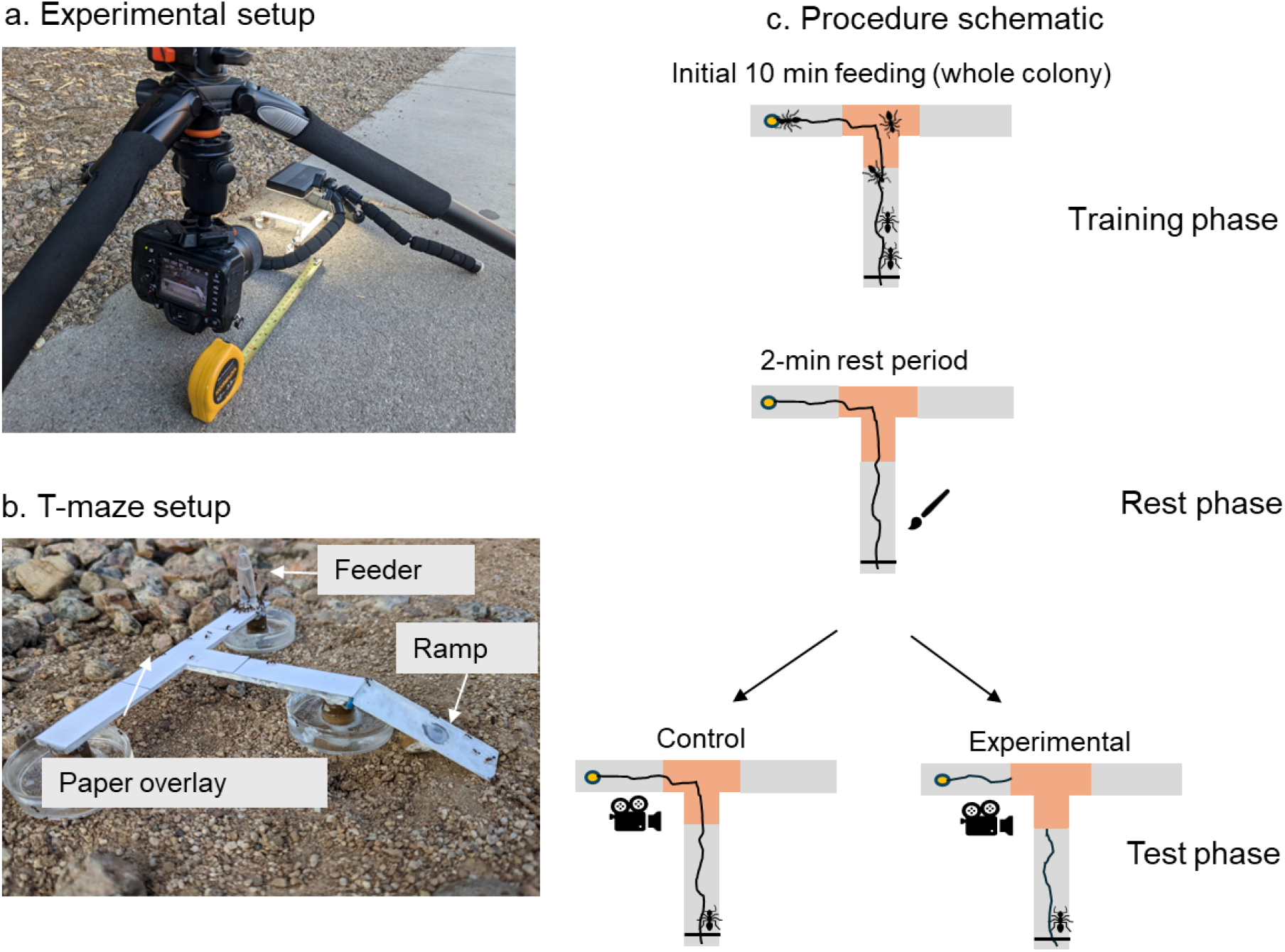
Experimental setup and procedure. (a) Camera and T-maze setup in the field, with measuring tape to determine the distance of the camera. (b) T-maze with feeder, ramp, and petri dish ‘moats’. (c) Schematic of the procedure. In the training phase, we allowed the whole colony to feed on the sucrose solution for 20 minutes in total (5 minutes near the colony entrance, 5 minutes on the T-maze stem, and 10 minutes on either arm of the T-maze that was selected randomly). We then brushed off the ants from the setup and feeder, and either applied the control treatment (removing and replacing the same paper section, highlighted with orange) or the experimental treatment (removal and replacement with fresh paper of the same section). Following that, we allowed one ant at a time to enter the setup, started a video recording, and recorded whether the ant made a choice or abandoned it. The video recording was stopped when the ant exited the setup (crossed the ‘line of commitment’which is shown in the figure above). The wiggly lines represent the pheromone trail. All the objects here (T-maze, ants, camera, brush, and pheromone trail) are schematics and not drawn as per the relative sizes of the objects in reality.

**Fig. 2.**
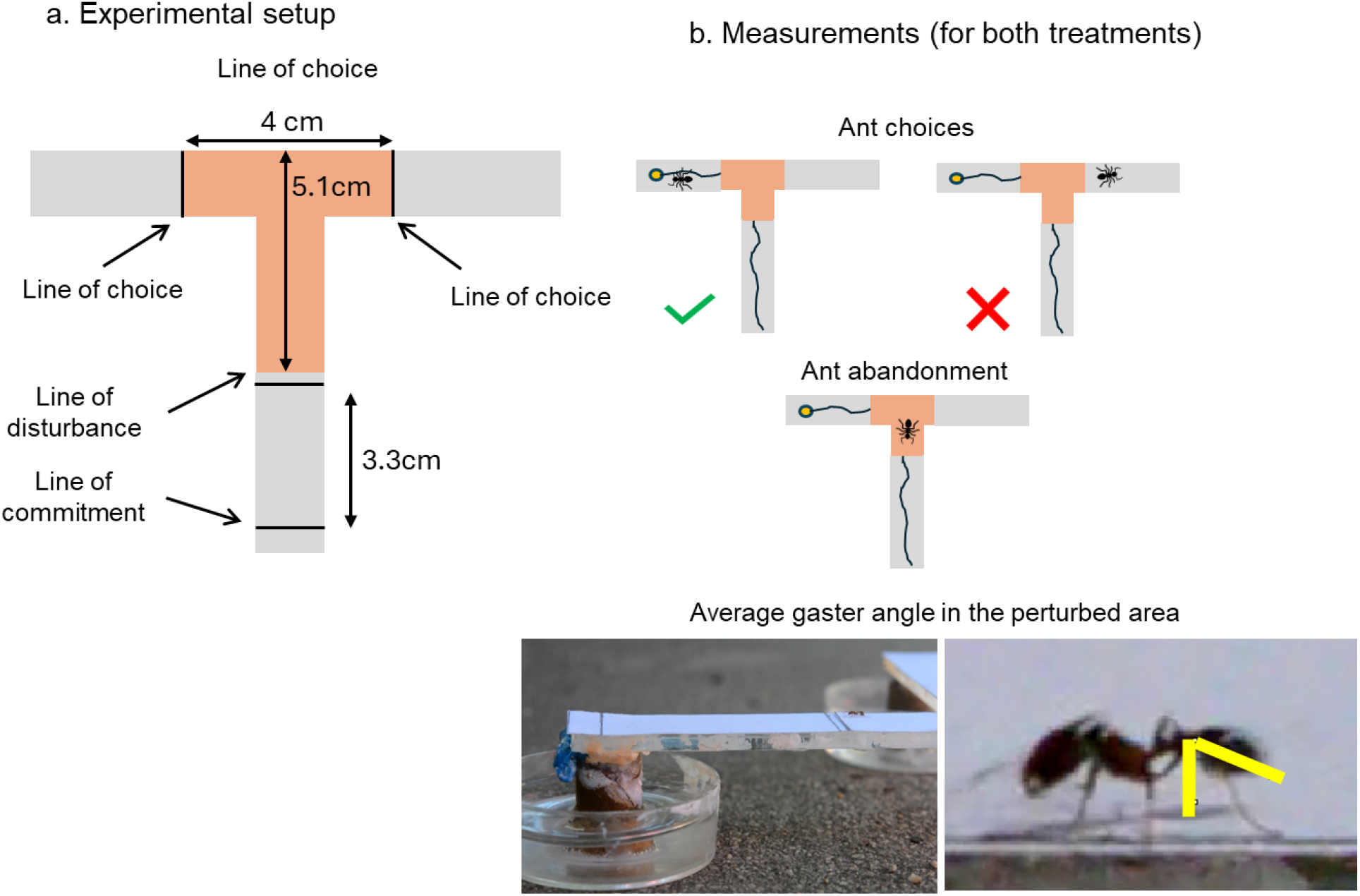
(a) Schematic of the pencil-drawn lines on the T-shaped paper cutout with measurements for each section. (b) Schematic of the measurements done during the experiments. We quantified three things for every ant: The first choice (left or right, i.e., correct or incorrect depending on feeder placement) or whether the ant abandoned the food source (turned back without crossing a choice line), and the average gaster angle on the return trip from a choice (while within orange area). The wiggly line represents the pheromone trail as present in the experimental treatment. Objects in Fig. 2b are not drawn to scale.

### Experimental procedure

We conducted all experiments between 9:30 am and 12 pm and 3:00 pm and 7:00 pm (July-November, 2022-25), when the ants were most active. We used two experiment types that only differed in the sucrose concentration used in the feeder (1.5 M (n = 116 trials) and 3 M (n = 86 trials); hereafter referred to as low sucrose and high sucrose, respectively). Both experiment types included trials in the control and experimental treatment (see ‘Test phase’ below). Each trial consisted of a training phase followed by a test phase.

### Training phase

We kept the T-maze ∼1m away from the colony’s nest entrance (the exact location near the nest and its position randomly chosen before every trial, Fig. 1a). We kept the feeder (filled with low or high sucrose solution) on the ground close to the colony entrance and allowed the ants to feed for 5 min since the first ant discovered and started feeding. We then moved the feeder onto the T-maze (with ramp access) for ants to access the feeder for another 5 min. Following that, we moved the feeder one last time to either of the T-maze arms randomly and allowed the colony to forage for another 10 min. This allowed the colony to lay a pheromone trail on the paper overlay towards the arm containing the feeder.

### Test phase

Following the 20-minute training phase, we removed the ramp and gently brushed off all the ants from the setup (including the feeder). Using nitrile gloves, we then removed the middle part of the T-shaped paper overlay (Length = 5.1 cm, Width = 4 cm) and replaced it in the same position (control treatment) or with a fresh paper section (experimental treatment). Thus, the control trials contained the same pheromone trail as laid by the ants in the training phase, but in experimental trials, there was a gap in the trail where the paper was replaced. We pointed the camera (Nikon D7000 DSLR) at a constant distance perpendicular to the setup so that it recorded the side-view of the majority of the T-maze stem (including the specific area where the paper had been replaced). Then, using the ramp, we allowed only a single ant to enter the setup. For every individual trial, once the focal ant entered the setup, we removed the ramp. We started the video recording once the ant crossed the ‘line of commitment’ (Fig. 2a). Once the ant returned and crossed the ‘line of commitment’ again, we stopped the recording. For the experimental treatment across both experiment types, we tested at most three ants in succession (each ant constituting an independent trial) using the same pheromone trail. This was possible because we put a fresh paper section on the T-maze for each experimental trial. However, for the control treatment, since we replaced the same paper section (with the trail on the paper), we tested just one ant per pheromone trail laid during the training phase. For both treatments, once the trial ended, we gently removed the ant using a paintbrush and placed it slightly away (>1m) from the colony to prevent it from attempting to return to the T-maze immediately.

### Data collection

We defined four relevant ‘lines’, drawn on the T-maze with pencil, to operationally assess the decisions made by foraging ants (Fig. 2a). The section of the paper, and thus the pheromone trail, removed in experimental trials was the one between the ‘line of disturbance’ and the two ‘choice’ lines. We defined an ant as having decided to forage once it crossed the ‘line of commitment’ and walked onto the maze (at which point we started recordings). We defined an ant as having made a choice when it crossed either the ‘line of choice’ towards the left or right arm of the T-maze. For each individual ant, we recorded (1) the first choice made (crossed either of the ‘line of choices’), (2) whether it abandoned the setup (turned back between the ‘line of choice’ and ‘line of disturbance’, i.e., never crossed a ‘line of choice’), and (3) the angle at which it held its gaster while walking between the ‘line of choice’ and the ‘line of disturbance’ on the way back from the top of the T-maze (extracted from the video). In ant species that use pheromone trail communication, workers typically bend/curve their gaster in a species-stereotypical fashion when laying a pheromone trail (Hangartner 1969; Hölldobler and Wilson 1990; Beckers et al. 1992; Jackson and Châline 2007). During this bending, the angle between the gaster and the vertical decreases compared to when the ant is not laying a trail (Wilson 1962; Beckers et al. 1992). Fire ant workers typically drag their gaster continuously on the ground (to lay pheromone trail) when they are returning to their nest from a food source (Wilson 1962; Hangartner 1969). We therefore used our measure of ‘gaster angle’ as a proxy for trail-laying behavior. We used Adobe Premiere Pro to extract all video frames in which the ant was between the ‘line of choice’ and the ‘line of disturbance’ on the T-maze, on the way back from the food source. Then we used ImageJ to quantify the gaster angle on every third frame. Fig. 1c illustrates the entire experimental protocol, and the Supplementary File 1 contains a more detailed explanation on the video analysis pipeline.

### Statistical analysis

We used Chi-square tests to compare the number of choices for either maze arm and the number of ants abandoning vs. staying on the setup between treatments. We used a binomial test to compare the number of ants choosing the left and right maze arms, and either abandoning or staying on the trail (i.e., making either choice) to a null expectation of random (50%) choice, within each treatment. We averaged the gaster angles across all frames measured for each ant (Range: 1-58 frames) for each trial. We then ran Kruskal-Wallis, Dunn posthoc, and Mann-Whitney U-Tests to compare these average gaster angles across ants making correct versus wrong choices between the control and experimental treatments. All data were analyzed in R (v. 4.5.2, R Core Team, 2024). Other than the base R, we used the following packages-*Dunn*.*test* (Dinno 2017), *gridExtra* (Auguie and Antonov 2016), *ggstats* (Larmarange 2023), *ggthemes* (Arnold et al. 2021), *rcompanion* (Mangiafico 2017), *and tidyverse* (Wickham et al. 2019). Detailed sample sizes for each experiment and its outcomes are also given in the Supplementary file S1 (Table S1 and S2).

### Data availability

The original data and the annotated R script. are made available in the Supplementary section (ESM_2.zip) and this GitHub repository: https://github.com/chjoshi24/Ch1-robustness-mechanisms-paper-2025. The detailed breakdown of year wise experimental data, information on datasheet columns, and video analysis pipeline can be found in Supplementary file 1.

## Results

In total, we tested 202 ants from 17 colonies (Range of ants tested per colony = 1 to 27) across four field seasons (2022-2025): 45 ants in the control treatment with the low sucrose feeder (35 with high sucrose), and 71 in the experimental treatment with the low sucrose feeder (51 with high sucrose; these numbers correspond to column totals in Fig. 4). Note that ants who abandoned are not included in the choice tests nor have gaster angles measured; the column totals in Fig. 3 are the sample sizes for just ants that made choices. A more detailed breakdown is provided in the Supplementary file S1 (Table S1 and S2).

**Fig. 3.**
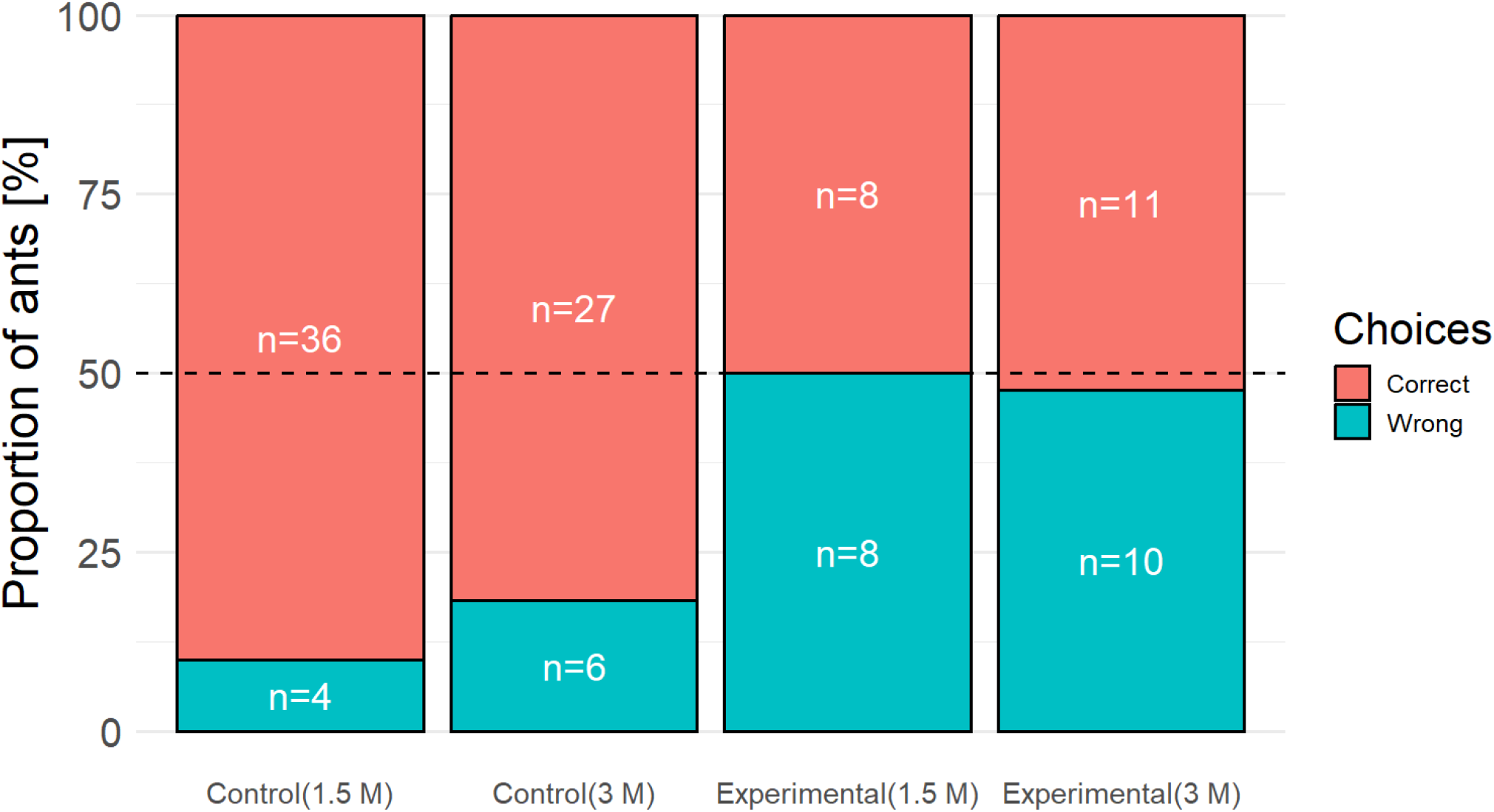
Proportion of ants making correct vs. wrong choices in control and experimental treatments across two experiment types. Ants made more correct choices in the control treatments (i.e., when the trail was intact; Binomial tests: low sucrose - p<0.001, n = 40; high sucrose - p<0.001, n = 33). The proportion of correct choices in either control or experimental treatments did not differ depending on food quality (Chi square tests: Control treatments- n = 73, χ^2^=1.02, df = 1, p = 0.31 and experimental treatments- n = 37, χ^2^ = 0.02, df=1, p = 0.89). The dotted line, at 50%, is the expectation if ants show no preference for either choice.

**Fig. 4.**
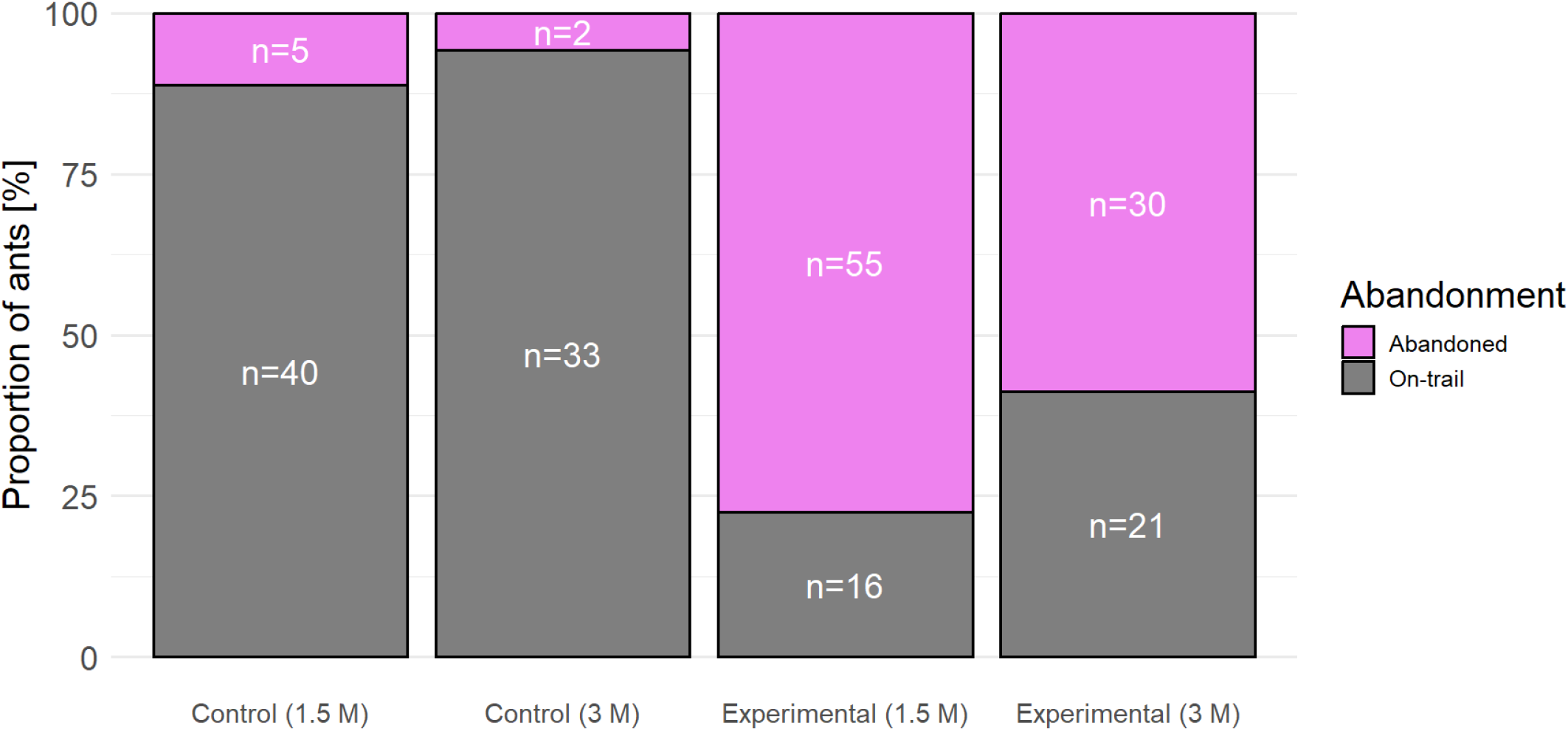
Proportions of ants that abandoned vs. stayed on the trail (correct and wrong choices combined). We defined all ants that turned and re-crossed the ‘Line of commitment’ without crossing the ‘Line of choice’ as ‘abandoning’. A significantly higher proportion of ants abandoned the food source in the experimental treatment compared to the control across both experiments (Chi-square tests: low sucrose - n = 116, χ^2^ = 48.57, df = 1, p<0.0001; high sucrose - n = 86, χ^2^ = 25.06, df = 1, p<0.0001); however significantly fewer abandoned in the high sucrose experimental treatment compared to the low sucrose one (Chi-square test: n = 122, χ2 = 4.88, df = 1, p = 0.03).

### Ant choices

Individually tested ants strongly preferred the arm of the T-maze that contained the food source, i.e., made a ‘correct’ choice (90% of ants in low sucrose treatment, 82% of ants in high sucrose treatment; Binomial tests: low sucrose - p<0.001, n = 40; high sucrose - p<0.001, n = 33; Fig. 3). However, when the pheromone trail was removed, the ants did not significantly prefer either direction (i.e., made ‘correct’ decisions at chance levels: 50% in low, 52% in high sucrose treatment; Binomial tests: low sucrose - p = 1, n = 16; high sucrose - p = 1, n = 21). In each condition (low vs high sucrose), the proportion of correct choices was higher in the control than the experimental (pheromone trail perturbed) treatment (Chi-square tests: low sucrose - n=40+16, χ^2^ = 10.86, df = 1, p<0.001; high sucrose - n = 33+21, χ^2^ = 5.33, df = 1, p = 0.02). We did not find an effect of food quality (sucrose concentration) on the proportion of correct choices for either control (Chi-square test: n = 40+33, χ^2^=1.02, df = 1, p = 0.31) or experimental treatments (Chi-square test: n = 16+21, χ^2^ = 0.02, df=1, p = 0.89).

### Ants abandoning the food source

In both low and high sucrose experiments, we found a significant relationship between the treatment and the proportion of ants that abandoned the resource, with more ants abandoning when the trail was perturbed (low sucrose: 10% in control, 77% in experiment; high sucrose: 6% in control, 59% in experiment; Chi-square tests: low sucrose - n = 45+71, χ^2^ = 48.57, df = 1, p<0.0001; high sucrose - n = 35+51, χ^2^ = 25.06, df = 1, p<0.0001; Fig. 4). There was no difference between low and high sucrose experiments in the control treatment (Chi-square test: n = 45+35, χ^2^ = 0.718, df = 1, p = 0.40). However, we found that abandonment was significantly lower in the high sucrose compared to the low sucrose experiments when the trail was removed (i.e., in the experimental treatment, Chi-square test: n = 71+51, χ2 = 4.88, df = 1, p = 0.03).

### Average gaster angle

Across both experiments, there were some ants that made a correct choice but did not feed on the sucrose, and some ants that first made a wrong choice but still fed on the sucrose (by moving across the other arm of the T maze to the feeder arm before exiting). For the analyses here, we excluded both types of ants, only including ants that indeed reached the food after a correct choice and fed, and those that, if they made an initial wrong choice, did not reach the food (and did not feed), in the gaster angle analysis. For completeness, we did analyze the gaster angles of the ants that were excluded in analyses here and included those results in Supplementary file S1 (Fig. S1 and S2). In addition to this, we did not quantify the gaster angles of the ants that abandoned the setup, since these ants failed to reach the food source.

We found that ants that made a correct choice had a significantly lower mean gaster angle compared to the ones that made a wrong choice in the high sucrose experiment (pooling control and experimental treatment - Kruskal-Wallis test: n=31, χ^2^ = 13.47, df = 2, p = 0.004; see Fig. 5 for the pairwise differences, Dunn’s post hoc test). This indicates that our gaster angle measure indeed quantifies trail laying, and that ants who reached a high-quality food source were more likely to lay a trail than those who did not. In the low-sucrose experiment, this was not the case (correctness of choice did not significantly affect gaster angle; however, there were only two incorrect choices in total, which means we could not clearly evaluate this question here - Fig. 5, Kruskal-Wallis test: n=36, χ2 = 2.21, df = 2, p = 0.331).

**Fig. 5.**
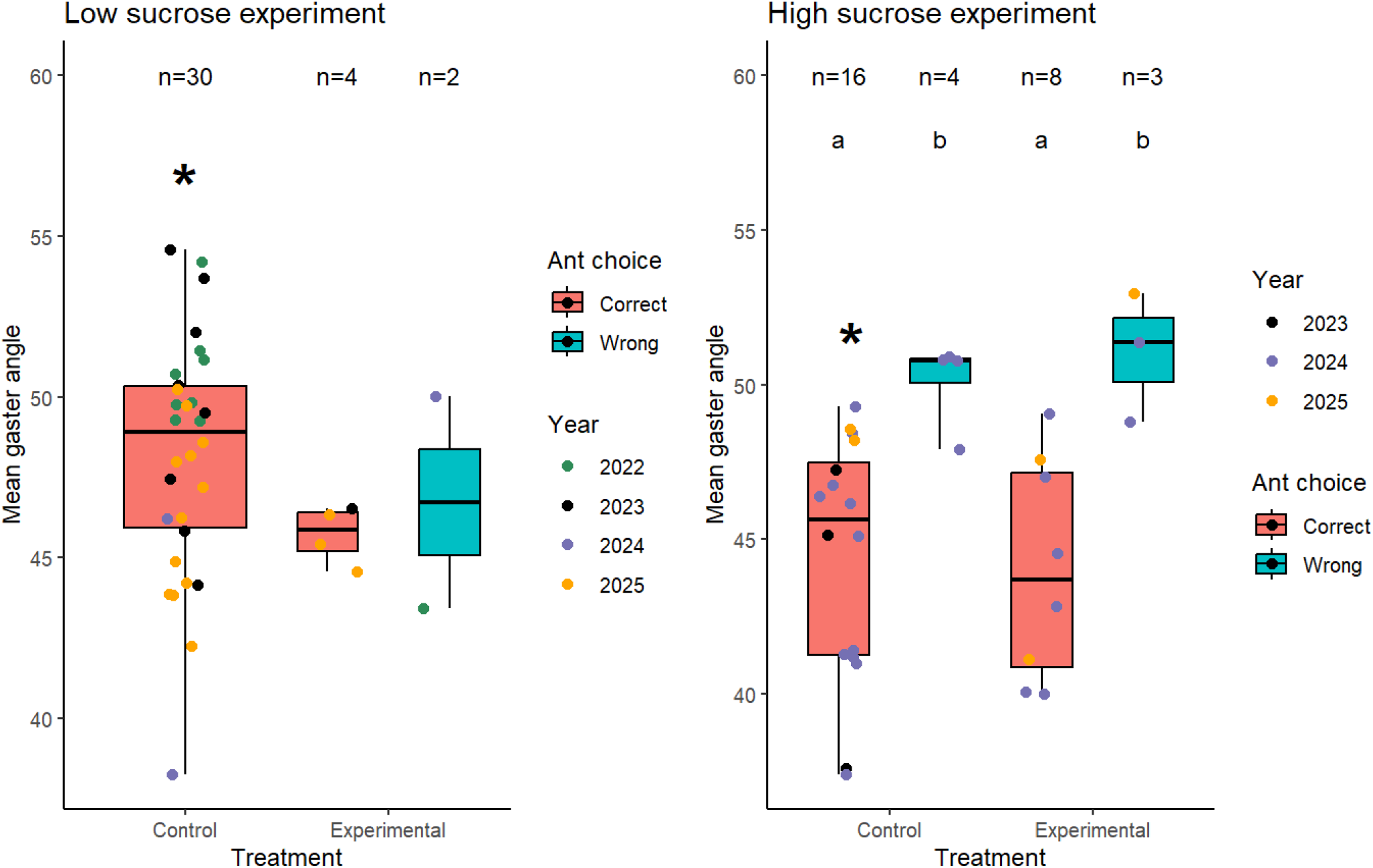
Gaster angle (averaged, for each ant, across video frames) in different treatments across low and high sucrose experiments. Only ants that made a correct choice and did feed on the feeder (red boxes), and ants that made a wrong choice and did not feed on the feeder (blue boxes), are included here. The year of the experiment is indicated by the color of the data points. We did not see a significant effect of treatment (control, with trail intact, or experiment, with trail removed). We also did not find a difference in gaster angles between ants that made correct or wrong choices in the low sucrose experiment. However, in the high sucrose experiment, ants that made a correct choice had a significantly lower gaster angle, i.e., were more likely to lay a new recruitment trail, compared to the ants that made a wrong choice. For the high sucrose experiment, the boxes that do not share the same letter are significantly different from each other (significant Kruskal-Wallis test followed by Dunn’s post hoc test). Additionally, in the control treatments across both experiments, the gaster angle was significantly lower for the high sucrose experiment compared to the low sucrose, but no such difference was found for the experimental treatments (Mann Whitney U-tests, differences indicated by the asterisks).

For the ants that made correct choices within control treatments across both experiments, the gaster angle was significantly lower in the high sucrose experiment compared to the low sucrose (difference between categories indicated by asterisk in Fig. 5, Mann-Whitney-U test: n = 30+16, W= 364, p = 0.004). The analogous difference was not significant for the experimental treatments, however again the sample size is fairly low (Fig. 5, Mann-Whitney-U test: n= 4+8, W = 20, p = 0.570).

## Discussion

We found that when confronted with an apparent dead-end in a pheromone trail, individual *S. xyloni* workers primarily reacted by turning back (Fig. 4). Those ants that continued towards the food source made random choices on the T-maze bifurcation in the absence of pheromone trail information (Fig. 3), suggesting that they were unable to use backup information sources such as their own memory or long-distance cues like vision or odor to locate the food source. When ants foraged on a more valuable food source, they remained more committed and did not turn around as often even in the absence of a pheromone trail; this was despite the fact that they still appeared to make random choices on the T-maze regardless of food source quality (Fig. 3). Our measure of gaster angle was lower when foragers were returning from high quality food rather than from an incorrect choice (and thus an unsuccessful trip) (Fig. 5). This confirms that the measure of gaster angle indeed indicates trail-laying. However, foragers equally reinforced the perturbed trail across both food qualities overall, and did not reinforce the trail more if it had disappeared (in control vs experiment).

Overall, our study thus showed a surprising lack of robustness to pheromone trail perturbation in fire ants. This failure indicates that individual *S. xyloni* may be highly reliant on the pheromone trail for navigating their way towards food sources, instead of using trails only to recruit naive ants (and then using their own memory to return to known food sources). Given that the pheromone trail serves as an externalized memory of the food source location, there may be low selection pressure for every forager to be successful at remembering the food source location (Feinerman and Traniello 2016). Studies from other fire ant species have shown that foragers can in principle use landmarks to navigate and memorize food locations, and pheromone trail presence can speed up this process (Stratton and Coleman 1973; Vander Meer et al. 2023). However, one of these studies shows that *Solenopsis saevissima* foragers took significantly longer to reach a food source in a multi-turn maze without a pheromone trail compared to when the trail was present, indicating that in that case the pheromone trail was also important for orientation during foraging (Stratton and Coleman 1973). It is also possible that *S. xyloni* foragers need more time to memorize the food source locations. All of our training phases only lasted 20 minutes. In the study mentioned above, authors also found that the *S. saevissima* foragers become less error-prone over 15-21 days in navigating their way towards the food source without using trails (Stratton and Coleman 1973). In addition to this, we only recorded the ant’s first choice in our experiment. So, it is possible that ants may eventually reach the feeder through exploration or during multiple exploratory trips at and around the previous location, even when a pheromone trail is perturbed. Nonetheless, this is likely to translate into a noisier, less reliable foraging process without pheromone communication.

*Solenopsis xyloni* and many other mass recruiting species (including other species from the *Solenopsis* genus) typically have big colony sizes with aggressive, but individually small workers (Valone and Kaspari 2005; Cerdá et al. 2013; Tschinkel 2013). This can enable colonies to monopolize a food source even against competing ants (Detrain and Deneubourg 2008; Lanan 2014). It may be that in a foraging system like this, food resources are only valuable to the colony when mass recruitment and strong symmetry breaking is possible (Lanan et al. 2012; Cerdá et al. 2013; Price et al. 2016), and colonies thus invest little in individual ability to reliably recover food sources. If that is the case, this would explain why foragers are so reliant on a social communication mechanism, the pheromone trail. Moreover, when the social information becomes outdated, e.g., the pheromone trail leads to a fully exploited or disappeared food source, ants have limited ways to counteract this information other than to let the trail evaporate (Czaczkes and Heinze 2015; Czaczkes et al. 2024). One of the ways in which ants may counteract this change is by upregulating trail laying towards another active food source nearby (Czaczkes and Heinze 2015), resulting in competition between the newly added trail and the outdated trail. However, this course correction is possible only for ants that have in fact located an alternative food source (Czaczkes and Heinze 2015) and even this strategy to update trail information does not always seem to be used (Czaczkes et al. 2024).

As noted above, the frequent abandonment of the food source by workers when the trail was perturbed might be interpreted as a lack of individual or collective robustness to this type of disturbance. In the robustness mechanism framework of Kitano 2004, we expected to find perhaps feedback (trail reinforcement or otherwise strengthened recruitment) or individual-level fail-safes (alternative information sources) in response to trail perturbation, but did not find either of these mechanisms in fire ants in our experiment. However, we could argue that an alternative interpretation to ‘absence of robustness’ is one of modularity or buffering. Ant colonies may maintain several concurrent, independent pheromone trails to different food sources, thereby constituting a modular system (Lanan 2014; and personal observations in *S. xyloni*). Disturbance of one of these trails likely does not affect foraging on other trails, thus containing the perturbation to one ‘module’ and maintaining (some) food intake by the colony. On the other hand, ants that ‘abandoned’ our T-maze may also not have abandoned foraging altogether; they may have switched to another available trail or resource. If individual workers have a choice between multiple maintained or discoverable resources, this would constitute a kind of ‘buffering’ mechanism in Kitano’s framework, where individual performance is only moderately impacted by the perturbation because of extra capacity. Future experiments are needed that involve tracking ants’ decisions to further understand the downstream consequences of individual-level and food-source-specific abandonment on colony-level foraging performance (in ants and in general across social insects).

Despite an overall high reliance on trail pheromone, we did find that even an apparently complete lack of information about which way the food was located did not prevent about 41% of foragers from attempting to find it when the resource they expected was of high quality. This illustrates a tradeoff between the risks of exploration (searching for a food source that may or may not be found) and the potential gains of exploitation (sucrose reward when food is found).

Overall, we found an apparent lack of ability to compensate for a disturbed pheromone trail in fire ants in our study, indicating that mass recruiting ant species like *S. xyloni* may not prioritize high robustness in this context. Possible reasons are a high reliance on strength in numbers and resulting low levels of individual reliability. This indicates a possible tradeoff between robustness, efficiency, and cost (Latty et al. 2011; Cook et al. 2014; Cabanes et al. 2015; Middleton and Latty 2016). Applying a general framework to categorizing robustness mechanisms allowed the identification of different possible approaches social insect colonies could take in dealing with perturbations of collective activity such as trail disruption. Application of such a framework may therefore be fruitful in a comparative context, by examining both different species (e.g., ones that differ in reliance on individual vs. mass foraging, or memory vs. communication) and examining different types of collective activity (e.g., foraging vs. task allocation vs. colony defense). Such endeavors may allow us to understand when and how robustness is prioritized and realized by collective systems of different types. Such studies will contribute to identifying the rules of life that are common to different scales of organization.

## Supporting information

R code and data files (updated R code)

Supplementary file 1 (Contents unchanged)

## Acknowledgments

We thank Michele Lanan for help with ant identification, Cycle Havel, Hayley Hananman, Tiffany Sessions, and Tyler Reese for contributions to data collection and video analysis, and the Staff at the Catalyst Studio, Main Library, University of Arizona, for help with T-maze laser cutting. We also want to thank Daniel Papaj, Judith Bronstein, and Tanya Latty, and the two anonymous reviewers for their valuable feedback on the manuscript. This work was supported by the Department of Ecology and Evolutionary Biology, University of Arizona, the National Science Foundation (Grant No. DBI 1564521 to Anna Dornhaus), and a project grant from the Graduate and Professional Student Council, University of Arizona to Chinmay Hemant Joshi.

