## Supplementary material for "Persist or Give up? Fire ants motivated to search for a high-quality food source even if they don’t know how to find it": R code and data files (updated R code): R code_robustness manuscript_2025.Rmd

````
---
title: "R code_robustness manuscript"
author: "Chinmay Joshi and Anna Dornhaus"
date: "2025-12-23"
output: html_document
---

```{r setup, include=FALSE}
knitr::opts_chunk$set(echo = TRUE)
```


IMP: Package installation step

If the packages below are not installed, this chunk of code will enable the user to install them before going ahead.
```{r}
pkgs <- c("tidyverse","ggplot2","ggthemes","ggstats",
          "gridExtra","dunn.test","rcompanion")
missing_pkgs <- setdiff(pkgs, rownames(installed.packages()))

if(length(missing_pkgs)) install.packages(missing_pkgs)
```


Required libraries/packages

```{r}
library(tidyverse) #For cleaning and reorganization of the data
library(ggplot2) #For plotting the graphs
library(ggthemes) #For formatting the graphs
library(ggstats)#For formatting the graphs
library(gridExtra) #For putting ggplots in a grid   
library(dunn.test) #For running Dunn's posthoc test
library(rcompanion) #For extracting significance letters from Dunn's posthoc test object using cldList function

```

CODE ORGANIZATION:
The code is organized as follows: Ant choice analysis, Ant abandonment analysis and gaster angle analysis. This is followed by the code for all the figures in the manuscript (Fig. 3, 4 and 5) and the supplementary section (Fig. S1 and S2). It is advisable to run the code in the order in which its written.


Importing the datasheets,and converting the relevant columns into factors.
Low sucrose experiment- 1.5 M sucrose
High sucrose experiment- 3 M sucrose
```{r}

low_sucrose_experiment_data<-read.csv("Combined ant choices_Low sucrose experiment.csv")

high_sucrose_experiment_data<-read.csv("Combined ant choices_High sucrose experiment.csv")

#Extracting the columns needed for choice and abandonment analysis
low_sucrose_experiment_choices<-data.frame(low_sucrose_experiment_data$Treatment,low_sucrose_experiment_data$Ant.choice)
high_sucrose_experiment_choices<-data.frame(high_sucrose_experiment_data$Treatment,high_sucrose_experiment_data$Ant.choice)

#Changing the column names
colnames(low_sucrose_experiment_choices)<-c("treatment","choices")
colnames(high_sucrose_experiment_choices)<-c("treatment","choices")

#Converting the columns into factors
low_sucrose_experiment_choices$treatment<-as.factor(low_sucrose_experiment_choices$treatment)
low_sucrose_experiment_choices$choices<-as.factor(low_sucrose_experiment_choices$choices)

high_sucrose_experiment_choices$treatment<-as.factor(high_sucrose_experiment_choices$treatment)
high_sucrose_experiment_choices$choices<-as.factor(high_sucrose_experiment_choices$choices)

#Creating labels specific to the experiment for the analysis below

#These two dataframes are used for the choice analyses below
low_sucrose_experiment_choices$label<-as.factor(ifelse(low_sucrose_experiment_choices$treatment=="Control","Control_1.5m","Exp_1.5m"))

high_sucrose_experiment_choices$label<-as.factor(ifelse(high_sucrose_experiment_choices$treatment=="Control","Control_3m","Exp_3m"))

#Ants stay on the trail when they are making a choice vs. when they abandon. So creating another column using the data above to indicate that. These two dataframes will be used for the abandonment analyses below

low_sucrose_experiment_abandonment<-low_sucrose_experiment_choices %>%
mutate(abandonment=ifelse(low_sucrose_experiment_choices$choices=="Abandoned","Abandoned","On-trail"))

low_sucrose_experiment_abandonment<-low_sucrose_experiment_abandonment[,c(1,3,4)]

high_sucrose_experiment_abandonment<-high_sucrose_experiment_choices %>% mutate(abandonment= ifelse(high_sucrose_experiment_choices$choices=="Abandoned","Abandoned","On-trail"))

high_sucrose_experiment_abandonment<-high_sucrose_experiment_abandonment[,c(1,3,4)]


```

ANT CHOICE ANALYSIS


Analyzing ant choices for low and high sucrose experiments
```{r}
#Filtering data where ants made either a correct or a wrong choice
low_sucrose_experiment_choices_only<-
  low_sucrose_experiment_choices %>% 
  filter(choices !="Abandoned")

high_sucrose_experiment_choices_only<-
  high_sucrose_experiment_choices %>%
  filter(choices !="Abandoned")

#Dropping the abandonment factor level
low_sucrose_experiment_choices_only$choices<-droplevels(low_sucrose_experiment_choices_only$choices)

high_sucrose_experiment_choices_only$choices<-droplevels(high_sucrose_experiment_choices_only$choices)

#Creating tables for the binomial and Chi-square tests 
low_sucrose_experiment_choices_only_table<-table(low_sucrose_experiment_choices_only[-3])

high_sucrose_experiment_choices_only_table<-table(high_sucrose_experiment_choices_only[-3])

#Binomial test for control treatments (Extracting relevant entries from the above choice tables)

binom.test(low_sucrose_experiment_choices_only_table[1],low_sucrose_experiment_choices_only_table[1]+low_sucrose_experiment_choices_only_table[3],0.5)

binom.test (high_sucrose_experiment_choices_only_table[1],high_sucrose_experiment_choices_only_table[1]+high_sucrose_experiment_choices_only_table[3],0.5)
#Significant!


#Binomial test for experimental treatments(Extracting relevant entries from the above choice tables) 
binom.test(low_sucrose_experiment_choices_only_table[2],low_sucrose_experiment_choices_only_table[2]+low_sucrose_experiment_choices_only_table[4],0.5)
#Not significant 

binom.test(high_sucrose_experiment_choices_only_table[2],high_sucrose_experiment_choices_only_table[2]+high_sucrose_experiment_choices_only_table[4],0.5)
#Non-significant 


#Chi-square tests to test whether proportion of correct choices are more in the control vs. experimental treatments across both experiments (Using the entire choice tables from above)

chisq.test(low_sucrose_experiment_choices_only_table,correct = F)
#Significant

chisq.test(high_sucrose_experiment_choices_only_table,correct = F)
#Significant

```


Comparing low and high sucrose experiments to examine whether the increased resource value resulted in changes in proportion of correct choices when the trail was perturbed.

Labels:
1.5m-Low sucrose experiment
3m- High sucrose experiment
```{r}
 
#Combining the choices from both experiments from the above data frames
all_exp_combined_data_choices<-rbind(low_sucrose_experiment_choices_only,high_sucrose_experiment_choices_only)

#Subsetting the data for each of the two treatments to compare between the experiments
all_exp_combined_control<-subset(all_exp_combined_data_choices,all_exp_combined_data_choices$treatment=="Control")

all_exp_combined_experimental<-subset(all_exp_combined_data_choices,all_exp_combined_data_choices$treatment=="Experimental")

#Dropping the unecessary factor levels for each dataframe
all_exp_combined_control$label<-droplevels(all_exp_combined_control$label)

all_exp_combined_experimental$label<-droplevels(all_exp_combined_experimental$label)

# Comparing proportion of correct choices for control and experimental treatments between the two experiments  
 
#Control treatment
all_exp_combined_control_choices_only<-data.frame(all_exp_combined_control$choices,all_exp_combined_control$label)

#Relabeling the columns and creating a table for the chi-sqaure test
colnames(all_exp_combined_control_choices_only)<-c("choices","exp_type")

all_exp_combined_control_choices_only_table<-table(all_exp_combined_control_choices_only)

#Chi-square test
chisq.test(all_exp_combined_control_choices_only_table,correct = F)
#Not significant

#Experimental treatment

all_exp_combined_experimental_choices_only<-data.frame(all_exp_combined_experimental$choices,all_exp_combined_experimental$label)

#Relabeling the columns and creating a table for chi-sqaure tests
colnames(all_exp_combined_experimental_choices_only)<-c("choices","exp_type")

all_exp_combined_experimental_choices_only_table<-table(all_exp_combined_experimental_choices_only)

chisq.test(all_exp_combined_experimental_choices_only_table,correct = F)
#Not significant

```

ANT ABANDONMENT ANALYSIS

Analyzing ant abandonment for low and high sucrose experiments
```{r}

#Creating tables for the binomial and Chi-square tests using the dataframes defined above 
low_sucrose_experiment_abandonment_table<-table(low_sucrose_experiment_abandonment[-2])
high_sucrose_experiment_abandonment_table<-table(high_sucrose_experiment_abandonment[-2])

#Chi-square tests on the above table
chisq.test(low_sucrose_experiment_abandonment_table,correct = F)
#Significant

chisq.test(high_sucrose_experiment_abandonment_table,correct = F)
#Significant

#Binomial test for control treatments (Extracting relevant entries from the above choice tables)
binom.test(low_sucrose_experiment_abandonment_table[1],low_sucrose_experiment_abandonment_table[1]+low_sucrose_experiment_abandonment_table[3],0.5)
#Significant difference

binom.test(high_sucrose_experiment_abandonment_table[1],high_sucrose_experiment_abandonment_table[1]+high_sucrose_experiment_abandonment_table[3],0.5)
#Significant difference

 
#Binomial test for experimental treatments(Extracting relevant entries from the above choice tables) 

binom.test(low_sucrose_experiment_abandonment_table[2],low_sucrose_experiment_abandonment_table[2]+low_sucrose_experiment_abandonment_table[4],0.5)
#Significant difference

binom.test(high_sucrose_experiment_abandonment_table[2],high_sucrose_experiment_abandonment_table[2]+high_sucrose_experiment_abandonment_table[4],0.5)
#Not significant 

```


Comparing abandonment across both experiments.
Labels:
1.5m-Low sucrose experiment
3m- High sucrose experiment

```{r}

#Combining abandonment data across both experiments
all_exp_combined_abandonment_only<-rbind(low_sucrose_experiment_abandonment,high_sucrose_experiment_abandonment)

#Control treatment comparison

all_exp_combined_control_abandonment_only<-all_exp_combined_abandonment_only %>% filter(treatment=="Control")

#Converting the column into a factor and removing uncessary factor levels
all_exp_combined_control_abandonment_only$abandonment<-as.factor(all_exp_combined_control_abandonment_only$abandonment)

all_exp_combined_control_abandonment_only$label<-droplevels(all_exp_combined_control_abandonment_only$label)

#Creating table for the chi-square test
all_exp_combined_control_abandonment_only_table<-table(all_exp_combined_control_abandonment_only[-1])

#Chi-square test
chisq.test(all_exp_combined_control_abandonment_only_table,correct = F)
#Not significant

#Experimental treatment comparison

all_exp_combined_experimental_abandonment_only<-all_exp_combined_abandonment_only %>% filter(treatment=="Experimental")

#Converting the column into a factor and removing uncessary factor levels
all_exp_combined_experimental_abandonment_only$abandonment<-as.factor(all_exp_combined_experimental_abandonment_only$abandonment)

all_exp_combined_experimental_abandonment_only$label<-droplevels(all_exp_combined_experimental_abandonment_only$label)

#Creating table for the chi-square test
all_exp_combined_experimental_abandonment_only_table<-table(all_exp_combined_experimental_abandonment_only[-1])

#Chi-square test
chisq.test(all_exp_combined_experimental_abandonment_only_table,correct = F)
#Significant!

```


GASTER ANGLE ANALYSIS


Loading the datasets and converting the relevant columns into numeric variables and factors
```{r}
#Low sucrose experiment
gaster_angle_low_sucrose<-read.csv("Combined gaster angles_Low sucrose experiment.csv")

#Converting the relevant columns to factors
gaster_angle_low_sucrose$Ant.choice<-as.factor(gaster_angle_low_sucrose$Ant.choice)
gaster_angle_low_sucrose$Treatment<-as.factor(gaster_angle_low_sucrose$Treatment)
gaster_angle_low_sucrose$Video<-as.factor(gaster_angle_low_sucrose$Video)
gaster_angle_low_sucrose$Year<-as.factor(gaster_angle_low_sucrose$Year)
gaster_angle_low_sucrose$Ant.fed.on.the.feeder<-as.factor(gaster_angle_low_sucrose$Ant.fed.on.the.feeder)

#Converting gaster angle measures to numeric variable type.
gaster_angle_low_sucrose$Angle.measurement.1<-as.numeric(gaster_angle_low_sucrose$Angle.measurement.1)
gaster_angle_low_sucrose$Angle.measurement.2<-as.numeric(gaster_angle_low_sucrose$Angle.measurement.2)
gaster_angle_low_sucrose$Angle.measurement.3<-as.numeric(gaster_angle_low_sucrose$Angle.measurement.3)
#Ignore the warnings here


#High sucrose experiment
gaster_angle_high_sucrose<-read.csv("Combined gaster angles_High sucrose experiment.csv")

#Converting the relevant columns to factors
gaster_angle_high_sucrose$Ant.choice<-as.factor(gaster_angle_high_sucrose$Ant.choice)
gaster_angle_high_sucrose$Treatment<-as.factor(gaster_angle_high_sucrose$Treatment)
gaster_angle_high_sucrose$Video<-as.factor(gaster_angle_high_sucrose$Video)
gaster_angle_high_sucrose$Year<-as.factor(gaster_angle_high_sucrose$Year)
gaster_angle_high_sucrose$Ant.fed.on.the.feeder<-as.factor(gaster_angle_high_sucrose$Ant.fed.on.the.feeder)


#Converting gaster angle measures to numeric variable type.
gaster_angle_high_sucrose$Angle.measurement.1<-as.numeric(gaster_angle_high_sucrose$Angle.measurement.1)
gaster_angle_high_sucrose$Angle.measurement.2<-as.numeric(gaster_angle_high_sucrose$Angle.measurement.2)
gaster_angle_high_sucrose$Angle.measurement.3<-as.numeric(gaster_angle_high_sucrose$Angle.measurement.3)
#Ignore the warnings here

```


Taking the averages of the gaster angle across a given frame in a video, and across multiple frames for a video.
```{r}

#Low sucrose experiment 
gaster_angle_low_sucrose_ready<-gaster_angle_low_sucrose %>%
  transmute(Video,Treatment,Ant.choice,Ant.fed.on.the.feeder, Year,mean_gaster_angle=rowMeans(select(.,Angle.measurement.1:Angle.measurement.3)))

#Reorganizing the data frame for the further analysis
gaster_angle_low_sucrose_ready<-as.data.frame(gaster_angle_low_sucrose_ready %>%
group_by(Video,Treatment,Ant.choice,Ant.fed.on.the.feeder,Year) %>%
summarise(mean_gaster_angle=mean(mean_gaster_angle,na.rm = T)))

#Removing NAs
gaster_angle_low_sucrose_ready<-na.omit(gaster_angle_low_sucrose_ready)

#High sucrose experiment
gaster_angle_high_sucrose_ready<-gaster_angle_high_sucrose %>%
  transmute(Video,Treatment,Ant.choice,Ant.fed.on.the.feeder,Year,mean_gaster_angle=rowMeans(select(.,Angle.measurement.1:Angle.measurement.3)))

#Reorganizing the data frame for the further analysis
gaster_angle_high_sucrose_ready<-as.data.frame(gaster_angle_high_sucrose_ready %>%group_by(Video,Treatment,Ant.choice,Ant.fed.on.the.feeder,Year) %>% summarise(mean_gaster_angle=mean(mean_gaster_angle,na.rm = T)))

#Removing NAs
gaster_angle_high_sucrose_ready<-na.omit(gaster_angle_high_sucrose_ready)

```


Further subsetting of the data: This data only contains the trials where ant made correct choices did feed on the sucrose, and if they made wrong choices, they did not feed. Following that running Kruskal-Wallis test followed by Dunn' post hoc test (if applicable).
```{r}
#Low sucrose experiment
gaster_angle_low_sucrose_ready_1<-gaster_angle_low_sucrose_ready %>% filter(!(Ant.choice=="Wrong" & Ant.fed.on.the.feeder=="Yes")) %>%
  filter(!(Ant.choice=="Correct" & Ant.fed.on.the.feeder=="No"))

gaster_angle_high_sucrose_ready_1<-gaster_angle_high_sucrose_ready %>% filter(!(Ant.choice=="Wrong" & Ant.fed.on.the.feeder=="Yes")) %>%
  filter(!(Ant.choice=="Correct" & Ant.fed.on.the.feeder=="No"))

#Comparing average gaster angles between the trials where ants made correct choices and wrong choices across control and experimental treatments

#Low sucrose experiment
#Kruskal test
kruskal.test(mean_gaster_angle ~ interaction(Treatment, Ant.choice), data = gaster_angle_low_sucrose_ready_1)
#Not significant

#Kruskal test
kruskal.test(mean_gaster_angle ~ interaction(Treatment, Ant.choice), data = gaster_angle_high_sucrose_ready_1)
#Significant difference

#Running Dunn's posthoc test for the high sucrose data to find the specific pairwise differences
dunn_test_high_sucrose_1<-dunn.test(gaster_angle_high_sucrose_ready_1$mean_gaster_angle,interaction(gaster_angle_high_sucrose_ready_1$Treatment,gaster_angle_high_sucrose_ready_1$Ant.choice))


```


Comparing average gaster angles: control-control and experimental-experimental across two experiment types
```{r}

gaster_angle_low_sucrose_ready_1$label<-as.factor(ifelse(gaster_angle_low_sucrose_ready_1$Treatment=="Control","Control_1.5m","Exp_1.5m"))

gaster_angle_high_sucrose_ready_1$label<-as.factor(ifelse(gaster_angle_high_sucrose_ready_1$Treatment=="Control","Control_3m","Exp_3m"))

combined_gaster_angle_data<-rbind(gaster_angle_low_sucrose_ready_1,gaster_angle_high_sucrose_ready_1)

  
combined_gaster_angle_data %>% filter(Treatment=="Control",Ant.choice=="Correct")%>%
  group_by(label)%>%
  summarise(mean(mean_gaster_angle))

wilcox.test(mean_gaster_angle ~ label, data = subset(combined_gaster_angle_data,Treatment=="Control" & Ant.choice=="Correct"))
#Significant

wilcox.test(mean_gaster_angle ~ label, data = subset(combined_gaster_angle_data,Treatment=="Experimental"& Ant.choice=="Correct"))
#Not significant

```


Comparing average gaster angles in the ants that made a correct choice vs. wrong choice across both treatments and whether they fed on the feeder or not (Whole data)
```{r}
#Low sucrose experiment
kruskal.test(mean_gaster_angle ~ interaction(Treatment, Ant.choice,Ant.fed.on.the.feeder), data = gaster_angle_low_sucrose_ready)
#Not Significant 

#High sucrose experiment
kruskal.test(mean_gaster_angle ~ interaction(Treatment, Ant.choice,Ant.fed.on.the.feeder), data = gaster_angle_high_sucrose_ready)
#Significant difference


#Running Dunn's posthoc test for the high sucrose data to find the specific pairwise differences
dunn_test_high_sucrose_2<-dunn.test(gaster_angle_high_sucrose_ready$mean_gaster_angle,interaction(gaster_angle_high_sucrose_ready$Treatment,gaster_angle_high_sucrose_ready$Ant.choice,gaster_angle_high_sucrose_ready$Ant.fed.on.the.feeder))

```

PLOTTING ALL THE GRAPHS 
 
Graphs as they appear in the main manuscript and supplementary section

Fig. 3: Stacked barplots of choices across both experiments
```{r}

#Calculating the proportion of correct vs. wrong choices for the graph 
choices_combined<-as.data.frame(all_exp_combined_data_choices %>%
  group_by(label,choices)%>%
  summarize(total=n())%>%
  group_by(label) %>%
  mutate(
    percent = total / sum(total) * 100
  ))

#Reordering the factor levels 
choices_combined$label<-factor(choices_combined$label,levels = c("Control_1.5m", "Control_3m", "Exp_1.5m", "Exp_3m"))

#Stacked Barplot
ggplot(choices_combined, aes(x = label, y = percent, fill = choices)) +
  geom_bar(stat = "identity", color = "black") +
  geom_text(
    aes(label = paste0("n=", total)),
    position = position_stack(vjust = 0.5),
    size = 6,           
    color = "white"
  ) +
  scale_y_continuous() +
  labs(x = "", y = "Proportion of ants [%]", fill = "Choices") +   
  theme_minimal(base_size = 14) +
  theme(
    axis.text.x = element_text(size = 13),
    axis.title.y = element_text(size = 22),
    axis.text.y = element_text(size = 18), 
    legend.title = element_text(size = 20),            
    panel.grid.major.x = element_blank()
  ) +geom_hline(yintercept = 50, linetype = "dashed")+
  scale_x_discrete(labels=c("Control_1.5m"="Control(1.5 M)","Control_3m"= "Control(3 M)", "Exp_1.5m"="Experimental(1.5 M)","Exp_3m" = "Experimental(3 M)"))+
  guides(fill=guide_legend(title="Choices"))

```


Fig. 4: Stacked barplots of abandonment across both experiments
```{r}

#Calculating the proportion of ants that abandoned vs. the ones that stayed one the trail (i.e., made a choice) for the graph 

abandonment_combined<-as.data.frame(all_exp_combined_abandonment_only %>%group_by(label,abandonment)%>%
  summarize(total=n()) %>%
    group_by(label) %>%
  mutate(
    percent = total / sum(total) * 100))

#Reordering the factor levels 
abandonment_combined$label<-factor(abandonment_combined$label,levels = c("Control_1.5m", "Control_3m", "Exp_1.5m", "Exp_3m"))

#Stacked Barplot
ggplot(abandonment_combined, aes(x = label, y = percent, fill = abandonment)) +
  geom_bar(stat = "identity", color = "black") +
  geom_text(
    aes(label = paste0("n=", total)),    
    position = position_stack(vjust = 0.5),   
    size = 6,                            
    color = "white"
  ) +
  scale_y_continuous() +
  labs(x = "", y = "Proportion of ants [%]") +
  theme_minimal(base_size = 14) +
  theme(
    axis.text.x  = element_text(hjust = 0.5, size = 13),  
    axis.text.y  = element_text(size = 16),               
    axis.title.y = element_text(size = 18),               
    legend.title = element_text(size = 18),              
    panel.grid.major.x = element_blank()
  ) +
  scale_x_discrete(labels = c(
    "Control_1.5m" = "Control (1.5 M)",
    "Control_3m"   = "Control (3 M)",
    "Exp_1.5m"     = "Experimental (1.5 M)",
    "Exp_3m"       = "Experimental (3 M)"
  )) +
  guides(fill = guide_legend(title = "Abandonment")) +
  scale_fill_manual(values = c("violet", "gray50"))

```


Fig. 5: Gaster angle Boxplots with Dunn posthoc letters (if applicable) and sample sizes, and year.

```{r}

#Defining color palette for the years
year_colors <- c("2022" = "seagreen","2023" = "black","2024" = "#7570b3","2025" = "orange")

#Low sucrose data
#Calculating treatment x ant choice sample sizes to put on the graph
low_sucrose_sample_size_1 <- gaster_angle_low_sucrose_ready_1 %>%
    group_by(Treatment, Ant.choice) %>%
    summarise(mean_angle = mean(mean_gaster_angle, na.rm = TRUE), n = n())

#This is to show the symbol for significance for the control treatment (Wilcoxon test above showed that gaster angles for the correct choices are signficantly different from each other between the control treatments across both experiments )
control_correct_symbol_low_sucrose <- gaster_angle_low_sucrose_ready_1 %>%
    filter(Treatment == "Control", Ant.choice == "Correct") %>%
    summarise(y = max(mean_gaster_angle, na.rm = TRUE) + 2) %>%
    mutate(Treatment = "Control",
           Ant.choice = factor("Correct", levels = levels(gaster_angle_low_sucrose_ready_1$Ant.choice)),
           label = "*")

#Boxplot
#Boxplot
boxplot_low_sucrose <- ggplot(gaster_angle_low_sucrose_ready_1,
  aes(x = Treatment, y = mean_gaster_angle, fill = Ant.choice)) +
    # Boxplots with increased spacing
    geom_boxplot(position = position_dodge(width = 1.2),
                 width = 0.6, color = "black",
                 outlier.shape = NA, coef = Inf) +
    # Jittered points that align perfectly with the boxplots
    geom_jitter(
      aes(color = Year, group = Ant.choice),
      position = position_jitterdodge(
        jitter.width = 0.40,
        jitter.height = 0,
        dodge.width = 1.0
      ),
      size = 2
    ) +
    # Sample size labels
    geom_text(data = low_sucrose_sample_size_1,
              aes(x = Treatment, y = 60,
                  label = paste0("n=", n),
                  group = Ant.choice),
              position = position_dodge(width = 1.0),
              inherit.aes = FALSE) +
    # Significance symbol for Control–Correct
    geom_text(data = control_correct_symbol_low_sucrose,
              aes(x = Treatment, y = y,
                  label = label,
                  group = Ant.choice),
              position = position_dodge2(width = 1.0,
                                         preserve = "single"),inherit.aes = FALSE,
              size = 10, fontface = "bold") +
  scale_color_manual(values = year_colors) +
  theme(panel.background = element_blank(),
          panel.grid.major = element_blank(),
          panel.grid.minor = element_blank(),
          panel.border     = element_blank(),
          axis.line        = element_line(color = "black")) +labs(title = "Low sucrose experiment",
         x = "Treatment",
         y = "Mean gaster angle",
         fill = "Ant choice")


#High sucrose experiment

#Extracting the letters from above Dunn'test object using cldList function from the rcompanion package. This is followed by renaming and reordering of the columns as per original data for correct placement of the letters in the graph

dunn_test_high_sucrose_letters_1 <- cldList(P.adjusted ~ comparisons, dunn_test_high_sucrose_1) %>%
    tidyr::separate(Group, into = c("Treatment","Ant.choice"), sep = "\\.") %>%dplyr::mutate(Treatment = factor(Treatment, levels = c("Control","Experimental")),
                  Ant.choice = factor(Ant.choice, levels = c("Correct","Wrong")))

#Calculating treatment x ant choice sample sizes to put on the graph
high_sucrose_sample_size_1 <- gaster_angle_high_sucrose_ready_1 %>%
    group_by(Treatment, Ant.choice) %>%
    summarise(mean_angle = mean(mean_gaster_angle, na.rm = TRUE), n = n())

#This is to show the symbol for significance for the control treatment (Wilcoxon test above showed that gaster angles for the correct choices are signficantly different from each other between the control treatments across both experiments )
control_correct_symbol_high_sucrose <- gaster_angle_high_sucrose_ready_1 %>%
    filter(Treatment == "Control", Ant.choice == "Correct") %>%
    summarise(y = max(mean_gaster_angle, na.rm = TRUE) + 2) %>%
    mutate(Treatment = "Control",
           Ant.choice = factor("Correct", levels = levels(gaster_angle_high_sucrose_ready_1$Ant.choice)),
           label = "*")

#Boxplot
boxplot_high_sucrose <- ggplot(gaster_angle_high_sucrose_ready_1,
aes(x = Treatment, y = mean_gaster_angle, fill = Ant.choice)) + 
  # Boxplots with increased spacing
  geom_boxplot(position = position_dodge(width = 1.0),
               width = 0.6, color = "black",
               outlier.shape = NA, coef = Inf) + 
  # Jittered points aligned perfectly with the boxplots
  geom_jitter(
    aes(color = Year, group = Ant.choice),
    position = position_jitterdodge(
      jitter.width = 0.40,
      jitter.height = 0,
      dodge.width = 1.0
    ),size = 2
  ) +labs(title = "High sucrose experiment", x = "Treatment",
y = "Mean gaster angle",fill = "Ant choice") +
  # Sample size labels
  geom_text(data = high_sucrose_sample_size_1,
            aes(x = Treatment, y = 60,
                label = paste0("n=", n),
                group = Ant.choice),
            position = position_dodge(width = 1.0),
            inherit.aes = FALSE) +
  # Dunn test letters
  geom_text(data = dunn_test_high_sucrose_letters_1,
            aes(x = Treatment, y = max(gaster_angle_high_sucrose_ready_1$mean_gaster_angle) + 5,
label = Letter,
                group = Ant.choice),
            position = position_dodge(width = 1.0),
            inherit.aes = FALSE) +
  # Significance symbol for Control–Correct
  geom_text(data = control_correct_symbol_high_sucrose,
            aes(x = Treatment, y = y,
                label = label,
                group = Ant.choice),
            position = position_dodge2(width = 1.0,preserve = "single"),
            inherit.aes = FALSE,
            size = 10, fontface = "bold", hjust = 2) +
  scale_color_manual(values = year_colors) +
  theme(panel.background = element_blank(),
        panel.grid.major = element_blank(),
        panel.grid.minor = element_blank(),
        panel.border     = element_blank(),
        axis.line        = element_line(color = "black"))

#Plotting the graphs side by side
grid.arrange(boxplot_low_sucrose, boxplot_high_sucrose, ncol = 2)

```


 


Fig S1 and S2: Boxplots of average gaster angles in the ants that made a correct choice vs. wrong choice across both treatments and whether they fed on the feeder or not across both experiment types(2 x 2 grids)

```{r}
#FigS1
#Calculating treatment x ant choice x ant fed or not sample sizes to put on the graph
low_sucrose_sample_size_2<-as.data.frame(gaster_angle_low_sucrose_ready %>%
  group_by(Treatment,Ant.choice, Ant.fed.on.the.feeder) %>%
  summarise(mean_angle = mean(mean_gaster_angle, na.rm = TRUE),
            n = n()))
  
ggplot(gaster_angle_low_sucrose_ready, aes(x = Treatment, 
               y = mean_gaster_angle, 
               fill = Ant.choice)) +
  geom_boxplot(position = position_dodge(width = 0.8))+  
  labs(title = "Low sucrose experiment",x = "Treatment",
       y = "Mean gaster angle",
       fill = "Ant choice") +geom_text(data =low_sucrose_sample_size_2, 
            aes(x = Treatment, 
                y =60,  # place labels above boxes
                label = paste0("n=", n),
                group = Treatment),
            position = position_dodge(width = 0.8),
            inherit.aes = FALSE)+geom_jitter(position = position_jitterdodge(jitter.width = 0.2, dodge.width = 0.8),
              alpha = 0.6, size = 2, color = "black")+facet_wrap(Ant.choice ~ Ant.fed.on.the.feeder, ncol = 2,labeller = labeller(Ant.fed.on.the.feeder = c("Yes" = "Fed on feeder", "No" = "Not fed")))+
 theme_minimal(base_size = 13) +
theme(
    panel.background = element_blank(),
    panel.grid.major = element_blank(),
    panel.grid.minor = element_blank(),
    panel.border     = element_blank(),
    axis.line        = element_line(color = "black"),
    strip.background = element_blank()
  )+scale_fill_manual(values = c("Correct" = "#A6CEE3", "Wrong" = "#B2DF8A"))

#FigS2
#Extracting the letters from above Dunn'test object using cldList function from the rcompanion package
dunn_test_high_sucrose_letters_2 <- cldList(P.adjusted ~ comparisons, dunn_test_high_sucrose_2)

#Renaming and reordering the columns as per original data for correct placement of the letters in the graph
dunn_test_high_sucrose_letters_2<- dunn_test_high_sucrose_letters_2 %>% 
  tidyr::separate(Group, into = c("Treatment","Ant.choice","Ant.fed.on.the.feeder"), sep = "\\.") %>%
  dplyr::mutate(
    Treatment = factor(Treatment, levels = c("Control","Experimental")),
    Ant.choice = factor(Ant.choice, levels = c("Correct","Wrong")),
    Ant.fed.on.the.feeder= factor(Ant.fed.on.the.feeder,levels= c("Yes","No"))
  )

#Calculating treatment x ant choice x ant fed or not sample sizes to put on the graph
high_sucrose_sample_size_2<-as.data.frame(gaster_angle_high_sucrose_ready %>%
  group_by(Treatment,Ant.choice, Ant.fed.on.the.feeder) %>%
  summarise(mean_angle = mean(mean_gaster_angle, na.rm = TRUE),
            n = n()))

#Boxplots with Dunn test letters and sample sizes

ggplot(gaster_angle_high_sucrose_ready, aes(x = Treatment, 
               y = mean_gaster_angle, 
               fill = Ant.choice)) +
  geom_boxplot(position = position_dodge(width = 0.8))+ labs(title = "High sucrose experiment",x = "Treatment",
       y = "Mean gaster angle",
       fill = "Ant choice") +geom_text(data =high_sucrose_sample_size_2, 
            aes(x = Treatment, 
                y =60,  # place labels above boxes
                label = paste0("n=", n),
                group = Treatment),
            position = position_dodge(width = 0.8),
            inherit.aes = FALSE)+geom_jitter(position = position_jitterdodge(jitter.width = 0.2, dodge.width = 0.8),
              alpha = 0.6, size = 2, color = "black")+facet_wrap(Ant.choice ~ Ant.fed.on.the.feeder, ncol = 2,labeller = labeller(Ant.fed.on.the.feeder = c("Yes" = "Fed on feeder", "No" = "Not fed")))+geom_text(data = dunn_test_high_sucrose_letters_2,
          aes(x = Treatment,
              y = max(gaster_angle_high_sucrose_ready$mean_gaster_angle) + 5,
              label = Letter,
              group = Ant.choice),
          position = position_dodge(width = 0.8),
          inherit.aes = FALSE)+theme(
    panel.background = element_blank(),
    panel.grid.major = element_blank(),
    panel.grid.minor = element_blank(),
    panel.border     = element_blank(),
    axis.line        = element_line(color = "black"),
    strip.background = element_blank()
  )+scale_fill_manual(values = c("Correct" = "#A6CEE3", "Wrong" = "#B2DF8A"))


```

END OF THE CODE 
````
