## Supplementary file 1 (Contents unchanged) for "Persist or Give up? Fire ants motivated to search for a high-quality food source even if they don’t know how to find it"

**Title: Persist or Give up? Fire ants motivated to search for a high-quality food source even if they don't know how to find it**

**Journal name:** Insect Sociaux

Chinmay Hemant Joshi [1*], and Anna Dornhaus [1]

[1] Department of Ecology & Evolutionary Biology, University of Arizona, Tucson, Arizona

**Supplementary results and discussion**

Across both low and high sucrose experiments, we had ants that either made a correct choice but did not feed on the sucrose, and ants that made a wrong choice but fed on the sucrose (by going to the food arm before exiting). We did not find a significant difference between the gaster angles regardless of the ant choice or its feeding status for the low sucrose experiment (Kruskal-Wallis test, n= 47, $\chi^2 = 9.48$, df = 6, p = 0.149, Fig. S1). However, for the high sucrose experiment, we found that the gaster angles were lower when the ants made a correct choice, compared to when they made a wrong choice (Kruskal-Wallis test, n= 50, $\chi^2 = 20.39$, df = 7, p = 0.005, Fig. S2). However, feeding status did not influence the gaster angle differences among the ants that made a correct or a wrong choice (Dunn's post hoc test, See Fig. S2 for the pairwise differences). This indicates that it may be sufficient for ants to decide to reinforce the trail while returning to the nest if they made a correct choice and encountered the feeder without feeding on it. Moreover, ants are most likely to reinforce the trail if they made a correct choice in the first place.

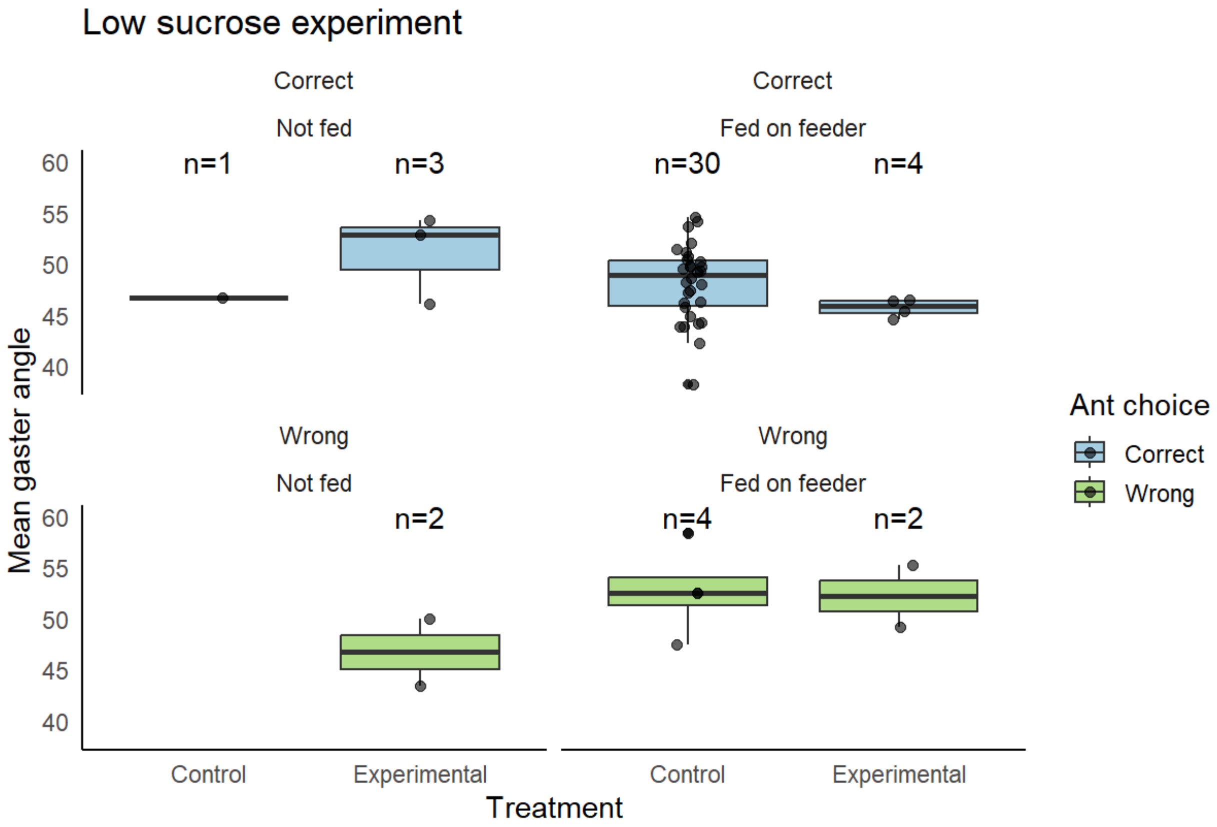


**Fig. S1** Mean gaster angles plotted against treatment type and faceted by whether ants made a correct choice or a wrong choice, and whether ants fed on the feeder or not. The sample size is indicated by the numbers on the top of each boxplot. We did not find a significant difference in the average gaster angles across treatments and regardless of the ant choice or whether the ants fed on the feeder or not (Kruskal-Wallis test: p = 0.149).

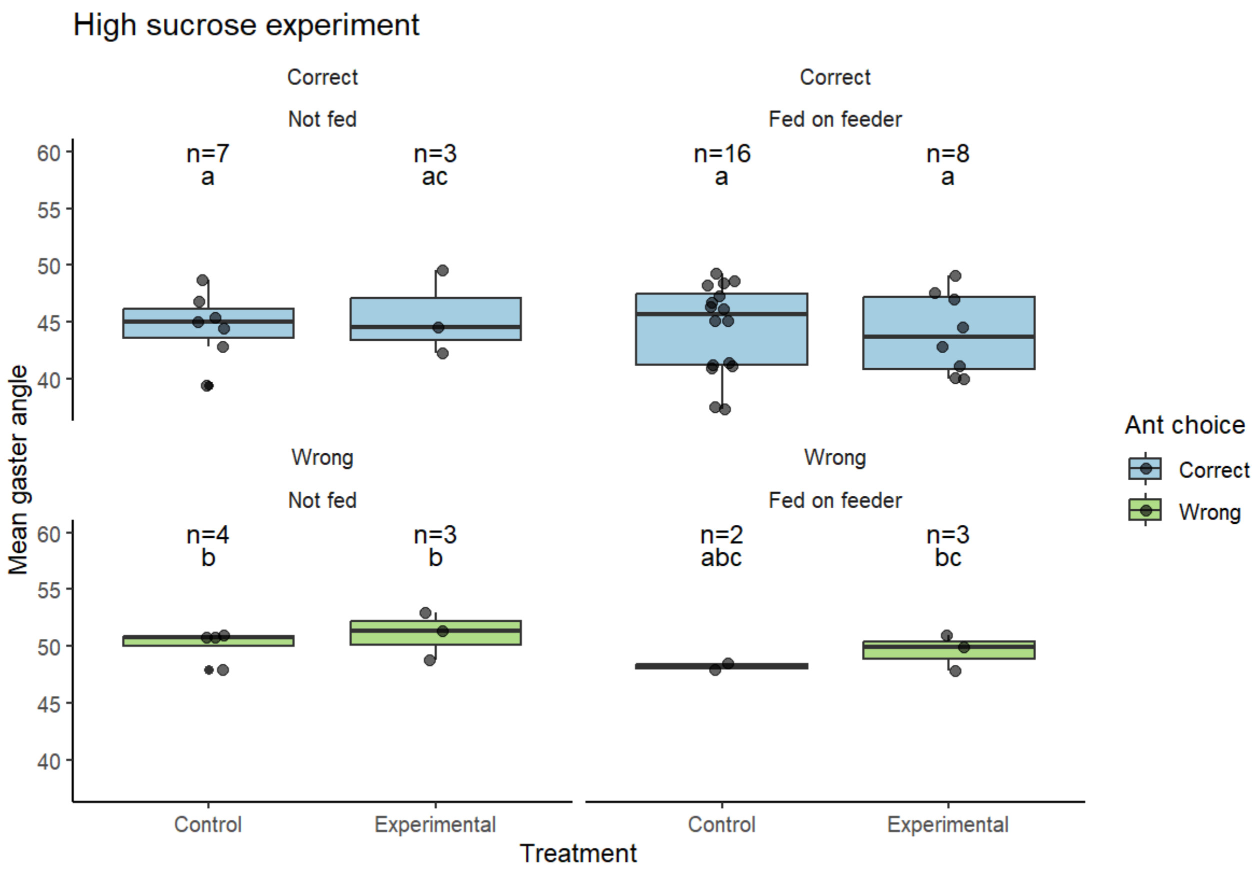


**Fig. S1** Mean gaster angles plotted against treatment type and faceted by whether ants made a correct choice or a wrong choice, and whether ants fed on the feeder or not. The sample size is indicated by the numbers on the top of each boxplot. Across all the eight plots, the boxes that do not share the same letters are significantly different from each other (Kruskal-Wallis test ($p = 0.005$), followed by Dunn's post hoc test). Overall, the average gaster angles are significantly lower when the ants made a correct choice, than when they made a wrong choice.

### Yearwise breakdown of the experimental data

**Table S1** Breakdown of choice and abandonment data for 1.5 M experiment for each year

| Treatment | Number of ants that made a correct choice | Number of ants that made a wrong choice | Number of ants that abandoned the setup | Total sample size |
|---|---|---|---|---|
| **2022 data (n= 40 total)** | | | | |
| Control | 9 | 3 | 1 | 13 |
| Experimental | 3 | 6 | 18 | 27 |
| **2023 data (n= 33 total)** | | | | |
| Control | 11 | 1 | 3 | 15 |
| Experimental | 1 | 0 | 17 | 18 |
| **2024 data (n= 10 total)** | | | | |
| Control | 3 | 0 | 1 | 4 |
| Experimental | 0 | 2 | 4 | 6 |
| **2025 data (n= 33 total)** | | | | |
| Control | 13 | 0 | 0 | 13 |
| Experimental | 4 | 0 | 16 | 20 |
| **Total sample size across all the years** | | | | 116 |

**Table S2** Breakdown of choice and abandonment data for the 3 M experiment for each year

| Treatment | Number of ants that made a correct choice | Number of ants that made a wrong choice | Number of ants that abandoned the setup | Total sample size |
|---|---|---|---|---|
| **2023 data (n= 12 total)** | | | | |
| Control | 4 | 1 | 0 | 5 |
| Experimental | 0 | 2 | 5 | 7 |
| **2024 data (n= 61 total)** | | | | |

| | | | | |
|---|---|---|---|---|
| Control | 21 | 5 | 2 | 28 |
| Experimental | 9 | 7 | 17 | 33 |
| **2025 data (n= 13 total)** | | | | |
| Control | 2 | 0 | 0 | 2 |
| Experimental | 2 | 1 | 8 | 11 |
| **Total sample size across all the years** | | | | 86 |

**Information on datasheet columns**

**Table S3** Combined ant choices Low and High sucrose experiment datasheets

| **Column** | **Information** |
|---|---|
| Date | This column recorded the date of data collection. |
| Time | This column recorded the time of data collection. |
| Observer | This column indicates the name of the experimenter who collected the data. |
| Colony ID | The ID of the colony is abbreviated as the colony number followed by the year of the data collection ('F' stands for Fall). It is important to note that a given colony was used only within that particular year. For example, C1F22 and C1F23 are different colonies that were used across 2022 and 2023 respectively. |
| Treatment | This column indicates where it was the control or experimental treatment. |
| Side/arm of the feeder | This column indicates the location of the feeder on the T-maze setup. The feeder was either placed on the right side or the left side on the setup. |
| Ant choice | This column indicates the first choice that the focal ant made. Ant could make one of the following three choices: Correct choice (going to the T-maze arm containing the feeder), Wrong choice (going to the T-maze arm not containing the feeder), OR abandoned (turn back without making any choice). |
| Ant fed on the feeder | This column indicates whether ant fed on the sucrose in the feeder. If the ant was on a feeder for more than 10 seconds continuously without moving, this column recorded 'Yes.' |
| Weather notes | This column recorded the weather (qualitatively) at the time of the experiment. |
| Surface temperature | This column recorded the temperature of the ground near the colony. It is important to note that we recorded this data from 2023 onwards. |
| Setup temperature | This column recorded the temperature of the paper overlay on the T-maze setup. It is important to note that we recorded this data from 2023 onwards. |

**Table S4** Combined gaster angles Low and High sucrose experiment datasheets

| **Column** | **Information** |
|---|---|
| Video | This column indicates the file name of the trial video. |
| Frame no. | This column indicates the number of the specific frame for the given video that was used for the gaster angle measurement. |

| | |
|---|---|
| Date of the video | This column recorded the date of data collection. |
| Colony ID | This column recorded the colony ID (for more information see under column information in Table S1). |
| Data of the analysis | This column recorded the date when the video was analyzed. |
| Treatment | This column indicates where it was the control or experimental treatment. |
| Year | This column recorded the year of data collection. |
| Observer doing the analysis | The observer who conducted the video analysis. It is important to note that the observers who collected the data were not always the ones who analyzed the videos. |
| Length in pixels of Zone2 | This column recorded the length of Zone 2 in the screenshot in pixels (See under 'Video analysis pipeline' below for more information) |
| Length in pixels of Zone1 | This column recorded the length of Zone 1 in the screenshot in pixels (See under 'Video analysis pipeline' below for more information) |
| End of bridge | This column indicates the location of the end of the T-maze stem in the video. This could appear on the right or the left side in the video. |
| Ant choice | This column indicates the first choice that the focal ant made. This column only has 'Correct' as the only option because the gaster angles were analyzed only for the ants that made a correct choice. |
| Angle measurement 1-3 | These three columns recorded the gaster angle measurements. Each row consists of 3 gaster angle measures for a given frame in a video. |
| Reason for NA (Angle measurement) | This column recorded why the observer could not analyze the particular frame. Some of the example responses include - 'Ant was too blurry', 'Ant not completely sideways' or 'Ant gaster not clearly visible.' |
| Gaster touching the paper | This column recorded whether the ant's gaster touched the paper overlay on the T-maze in the given frame. There were three possible responses here: 'Yes' (The gaster touched the paper), 'No' (The gaster did not touch the paper), OR 'NA' (The observer could not tell whether the gaster touched the paper or not). |

**Video analysis pipeline**

We quantified three key measures: Proportion of ants making correct choices, proportion of ants abandoning the food source, and the average gaster angle. The first two measures were recorded by the observers during the experiment directly in the datasheet. The average gaster angle was quantified from analyzing the videos taken during the experiment. The experiment videos captured the side-view of the ant moving on the T-maze stem (Fig. S1). Fire ants extrude their stinger and drag it continuously to lay pheromone trail on the surface while returning from a food source. So, we used gaster angle as a proxy for the pheromone trail laying- lower gaster angle indicates higher pheromone trail reinforcement.

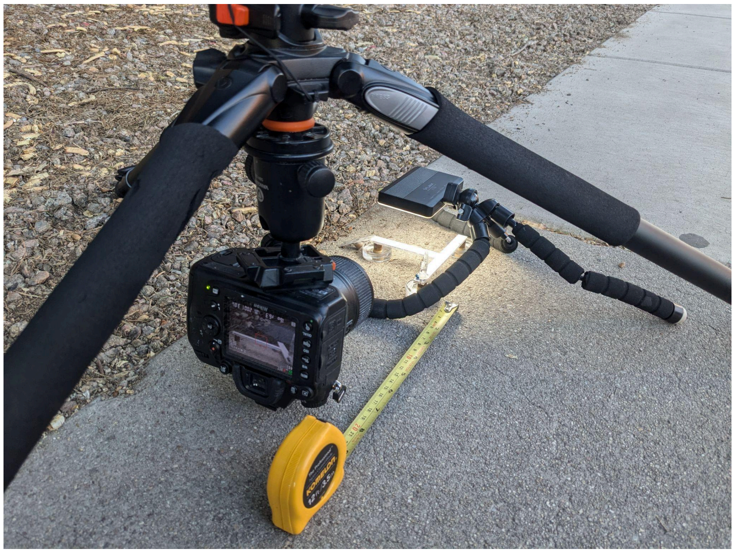

**Fig. S1** Experimental setup in the field

**Quantifying the average gaster angle from the videos**

We quantified the gaster angles only for all the trials where the ants either made a correct choice or a wrong choice. As mentioned in the main text, we defined relevant ‘lines’ by lightly pencil drawing them on the T-shaped paper overlay to operationally assess the ant decisions (Fig. S2). These lines allowed us to divide the side view of the paper overlay into three distinct zones- Zone 1, Zone 2 and Zone 3 (Fig. S3). We measured the gaster angles specifically in Zone 1, because this was the area where either the control treatment (removal and replacement of the paper containing the pheromone) or the experimental treatment (removal of the trail) was applied.

While taking the experimental videos in the field, we used slightly different zoom settings across the years, due to which only the part of Zone 1 on the T-maze stem was in the camera focus. So, for a given year, we fixed the effective Zone 1 edge length in centimeters (which was visible in the video), and obtained its length in pixels using Zone 2 length using Image J. Using this Zone 1 length in pixels, we drew a rectangle to define the effective Zone 1. Once the effective Zone 1 was clearly defined, we used Adobe Premier Pro to divide the video (24 frames per second) into individual frames, and extracted frame screenshots every 3 frames for the period where the ant was within this effective Zone 1, before the ant crossed into Zone 2. An ant’s gaster is equivalent to an ellipsoid (or an ellipse in two dimensions), and to measure the gaster angle against the vertical, one would need to define three points- two points on the ellipse circumference, and one point outside the ellipse defining the vertical (Fig. S4). So, during gaster angle quantification, the uncertainty in the measures could be introduced by the choice of either of these three points. So, using this operational definition, we quantified three measures of gaster using the ‘Angle tool’ in ImageJ. Then using R, we first averaged across these three gaster angle measurements for a given frame, and then averaged the measures across all the frames to get the final measure for the average gaster angle for an ant in a given trial.

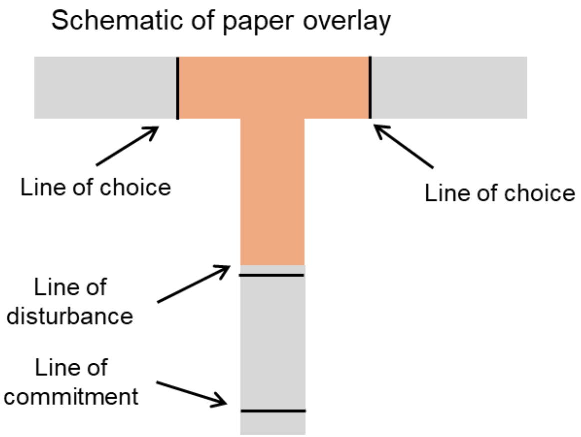


**Fig. S2** Schematic of the relevant pencil drawn lines on the T-shaped paper overlay (See under 'Data collection' in the main manuscript for more information on the lines).

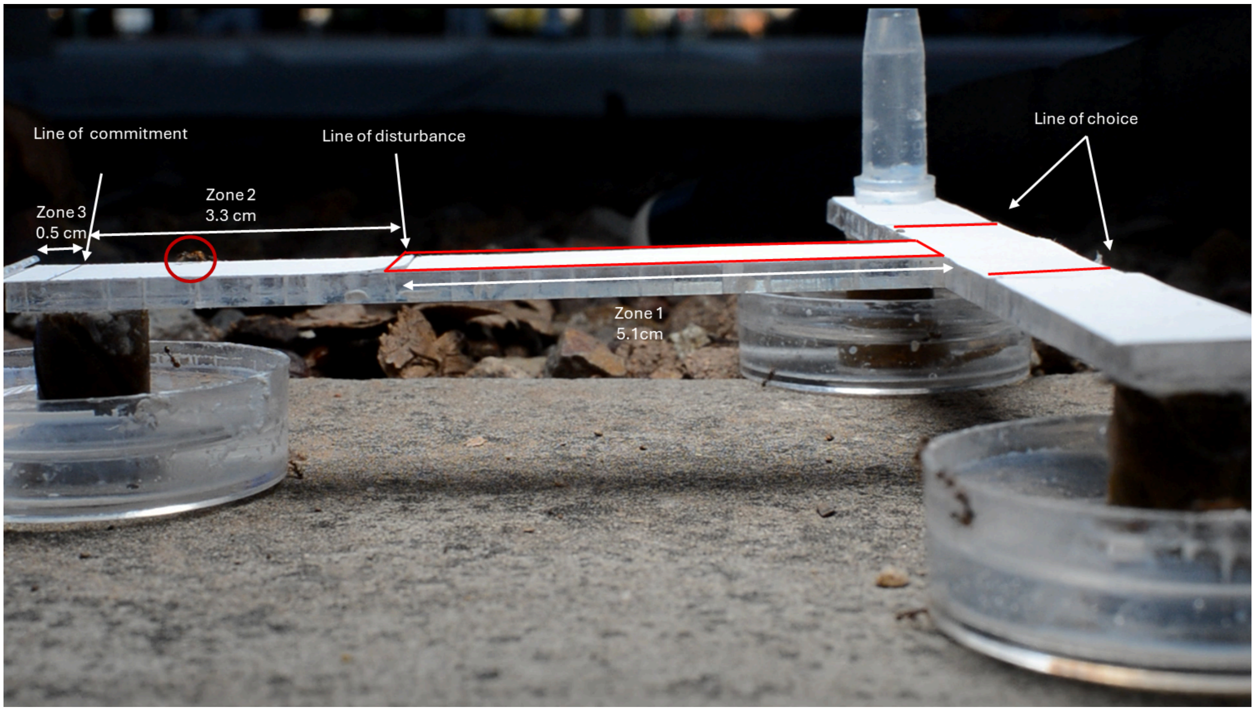


**Fig. S3** Example screenshot to illustrate the three zones. Zone 3 is between the edge of the T-maze and the foraging line. Zone 2 is between the line of commitment and the line of disturbance. Zone 1 is between the line of disturbance and lines of choice. Since only a part of the Zone 1 was in camera focus, all the gaster angle measurements were done in the area indicated by the red rectangle.

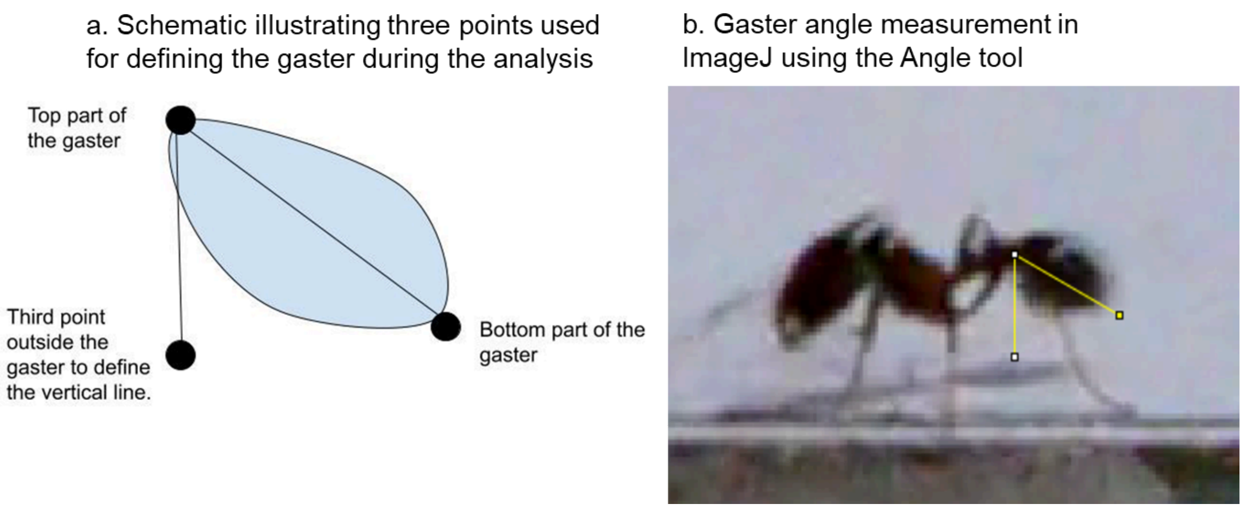


**Fig. S4** (a) Schematic illustrating three points used for measuring gaster angles (b) Example screenshot illustrating the gaster angle measurement using the angle tool on Image J.
